# Multiple conformational states assembly of multidomain proteins using evolutionary algorithm based on structural analogues and sequential homologues

**DOI:** 10.1101/2023.01.15.524086

**Authors:** Chunxiang Peng, Xiaogen Zhou, Jun Liu, Minghua Hou, Stan Z. Li, Guijun Zhang

**Author notes:** Correspondence should be addressed to Guijun Zhang.

## Abstract

With the breakthrough of AlphaFold2, nearly all single-domain protein structures can be built at experimental resolution. However, accurate modelling of full-chain structures of multidomain proteins, particularly all relevant conformations for those with multiple states remain challenging. In this study, we develop a multidomain protein assembly method, M-SADA, for assembling multiple conformational states. In M-SADA, a multiple population-based evolutionary algorithm is proposed to sample multiple conformational states under the guidance of multiple energy functions constructed by combining homologous and analogous templates with inter-domain distances predicted by deep learning. On a developed benchmark dataset containing 72 multidomain proteins with multiple conformational states, the performance of M-SADA is significantly better than that of AlphaFold2 on multiple conformational states modelling, where 29/72 (40.3%) of proteins can be assembled with a TM-score >0.90 for highly distinct conformational states with M-SADA while AlphaFold2 does so in only 2/72 (2.8%) of proteins. Furthermore, M-SADA is tested on a developed benchmark dataset containing 296 multidomain proteins with single conformational state, and results show that the average TM-score of M-SADA on the best models is 0.913, which is 5.2% higher than that of AlphaFold2 models (0.868).

---

Much progress has been made in protein structure prediction as a result of decades of effort^1^. In particular, the end-to-end sequence-to-structure training approaches, such as AlphaFold2^2^, built on the attention and equivariant transformer networks, have achieved unprecedented modelling accuracy in the protein structure prediction as witnessed in the CASP14 experiment^3, 4^. AlphaFold2 predicts the structures of many protein targets at or near experimental resolution, including some multidomain proteins^2, 5^. Accordingly, the DeepMind and the European Molecular Biology Laboratory (EMBL)-European Bioinformatics Institute collaborated to create a new data resource, AlphaFold Protein Structure Database (AlphaFold DB)^6^, which has enabled an unprecedented expansion of the structural coverage of the known protein-sequence space. However, AlphaFold2 is hardly optimized specifically for multidomain proteins, and the confidence score of the AlphaFold2 model is highly correlated with whether a target has homologues in the Protein Data Bank (PDB) ^7^. Thus, the structure prediction of multidomain proteins remains relatively unreliable when the number of homologous templates or sequences of related proteins is insufficient for inferencing relative domain orientations^8^. Another important limitation is that the predicted structural models do not provide insight into conformational dynamics. AlphaFold2 predictions typically generate one of the states of the protein, but understanding the mechanism of action will require knowledge of the broader conformational landscape^9^. This is also true for multidomain proteins, for which function is intimately connected to changes in tertiary and quaternary structure spanning multiple conformational states. The development of computational methods to address the gap between single and multiple domains, and that between single and multiple conformational states can yield great dividends^9^.

In general, domain assembly methods are divided into two major categories: *ab initio* methods and template-based approaches. *Ab initio* methods can be adopted to predict multidomain protein full-chain structures when reliable structures are available for individual domains, such as Rosetta^10^, AIDA^11^ and GalaxyDomDock^8^. However, *ab initio* methods may leave the domain structures largely randomly oriented in the final model^12^. Template-based methods use template information to infer domain orientations, making the orientation between domains more reliable to a certain extent. Representative template-based domain assembly methods include DEMO^12, 13^ and SADA^14^. In our developed DEMO, docking-based domain assembly simulations are performed to generate full-chain models of multidomain proteins. Based on DEMO, a domain enhanced modelling method using cryo-EM (DEMO-EM) is proposed^15^, to create accurate full-chain structural models for multidomain proteins from cryo-EM density maps. In our developed SADA^14^, analogous full-chain templates are identified through domain-level structure alignments, and the optimal positions and orientations of all domains are simultaneously searched to generate the full-chain model through a two-stage differential evolution algorithm guided by the energy function, with an inter-residue distance potential predicted by deep learning. However, SADA ignores the importance of available homologous templates for inferring inter-domain orientations.

Although homologous templates that can be used to model the full-chain structures of multidomain proteins are frequently unavailable due to the limited structures in PDB^11^, the high-quality models in AlphaFold DB may be an effective complement to PDB in modelling multidomain protein structures. Therefore, the adequate use of sequential homologous and structurally analogous templates may be an important approach for modelling multidomain proteins. In fact, many multidomain proteins are expected to undergo conformational changes involving domain orientations to form different conformational states^8^. For example, the balance of a kinase’s active and inactive state must be regulated precisely in a cell. In an oversimplified picture, only these two highly distinct conformations exist for a kinase. While the vast majority of ATP competitive inhibitors binds to the active conformation of kinases, and a few small molecules (e.g. the anticancer drug imatinib) bind selectively to the inactive form of ABL1^16^. For advanced protein structure prediction methods, such as AlphaFold2, although they can accurately predict backbone and side chain conformations, they are for a particular conformational state^16^.

In this work, a method called M-SADA is proposed to model multiple conformational states of multidomain proteins, where potential domain orientations are explored through a multiple population-based evolutionary algorithm on the basis of sequential homologues, structural analogues and deep learning predicted inter-domain distance. On a development multidomain protein dataset with multiple conformational states, M-SADA achieves an average TM-score of 0.87, which is 16.0% higher than that of AlphaFold2 (0.75). Meanwhile, the results on large-scale benchmark datasets with single conformational state proteins indicate that M-SADA is significantly better than state-of-the-art domain assembly methods, such as SADA, and full-chain modelling methods, such as AlphaFold2.

## Results

### M-SADA overview

The pipeline of M-SADA is shown in **Figure 1**. Starting from the input full-chain sequence and computationally predicted (or experimentally solved) domain structures, the full-chain structural analogues are detected from our developed multidomain protein structure database (MPDB) according to the input protein domain models^14, 17^. Then, the homologous templates are successively searched from MPDB, PDB, AlphaFold DB90 and AlphaFold DB according to the input full-chain sequence, where AlphaFold DB90 represents the models with an average (per-residue local distance difference test) pLDDT ≥90 in AlphaFold DB. Subsequently, the analogous and homologous templates are used to design multiple energy functions, which include different template restraints, physical constraints and residue distance information predicted by the in-house inter-domain distance prediction method DeepIDDP^18^. A multiple population-based evolutionary algorithm is designed to explore and exploit the domain orientations based on multiple energy functions. Finally, full-chain models from different populations are selected and ranked using our developed model quality assessment method DeepUMQA2^19^.

**Figure 1.**
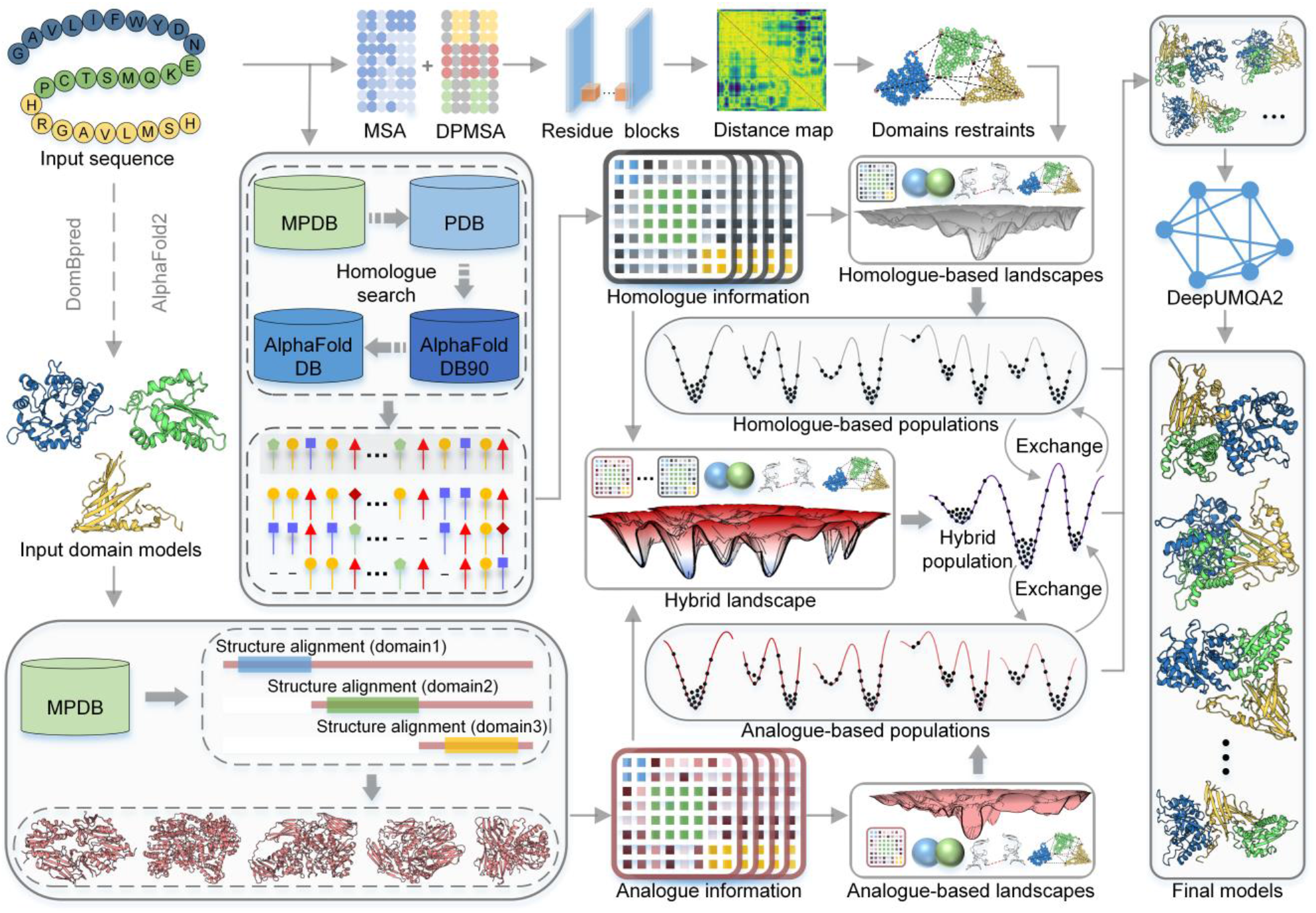
Pipeline of M-SADA.

### Datasets

To examine the ability of M-SADA to model multidomain proteins with multiple conformational states, its effectiveness of M-SADA to model multidomain proteins with single conformational state, and its domain assembly performance, we tested and discussed M-SADA on three different test sets. In order to study the ability of M-SADA on modelling multidomain proteins with multiple conformational states, M-SADA is tested on 72 multidomain proteins with multiple conformational states. The benchmark dataset is constructed in accordance with the following criteria: (1) all multidomain proteins in PDB are clustered at 100% sequence identity cut-off, and the TM-scores between different conformations in each cluster (i.e. each multidomain protein) are calculated, (2) the multidomain proteins with at least one pair of conformations with TM-score ≤0.75 are selected, (3) the two structures with the largest structural difference in the multidomain proteins are selected as the representative conformational states of the multidomain protein, (4) these multidomain proteins are removed if residues are missing at the domain boundary for representative conformations and (5) the final benchmark set with multiple conformational states (containing 72 multidomain proteins) is generated with a 40% sequence identity cut-off. To the best of our knowledge, this multidomain protein dataset is the first to be constructed systematically to investigate the ability to model multidomain protein structures with multiple conformational states.

In order to test the effectiveness of M-SADA, we reassemble 296 single conformational state multidomain proteins randomly selected from AlphaFold DB. The selection criteria for the 296 proteins are as follows: (1) 90% of the residues are solved in the native structure, (2) the sequence identity to the training set of DeepIDDP is less than 40% and (3) the sequence identity between each other is less than 40%.

In order to fairly compare the domain assembly performance of M-SADA with other domain assembly methods at the same level, all 356 multidomain proteins used in the SADA benchmark dataset are selected to test the performance of M-SADA^14^. The sequence identity amongst all the 356 proteins is less than 30%. This benchmark includes 166 2-domain (2dom), 69 3-domain (3dom), 40 ≥4-domain (m4dom) and 81 discontinuous domain (2dis) proteins. The maximum number of domains in m4dom is 7.

### Modelling multidomain proteins with multiple conformational states

Domains between multidomain proteins frequently interact to perform higher-level functions. Therefore, the conformational states of some multidomain proteins are often not unique. These different conformational states are critical for further studies on protein function and drug design. However, currently available protein structure modelling methods do not seem to specifically address this issue. In this section, we investigate whether M-SADA exhibits the ability to model the full-chain structures of 72 multidomain proteins with different conformational states, and then compare it with state-of-the-art method AlphaFold2. Here, the structure models of AlphaFold2 in each multidomain protein are predicted using its latest standalone package. The individual domain models for M-SADA assembly are from the first model of AlphaFold2, and 100% sequence identity cut-off is used to exclude templates.

To investigate whether the output model contains multiple conformational states, all assembled (or predicted) models for each protein in the same multidomain protein cluster are used to calculate the TM-score to the native structures, and then the maximum TM-score is taken. The results are summarized in **Supplementary Table S1** and the details for each multidomain protein cluster are shown in **Supplementary Table S2**. For all conformations of the 72 multidomain proteins, the average TM-score of AlphaFold2 is 0.77 while the average TM-score of M-SADA is 0.87, which is 13.0% higher than that of AlphaFold2. However, some multidomain proteins may contain similar conformational states, which affects the analysis of the multiple conformational states modelling ability to a certain extent. Therefore, we focus on the two conformational states with the maximum structural difference for each multidomain protein cluster. For the state 1 of the 72 multidomain proteins, the average TM-score of M-SADA is 0.88, which is 17.3% higher than that of AlphaFold2 (0.75). For the state 2 of the 72 multidomain proteins, the average TM-score of M-SADA is 0.86, which is 13.2% higher than that of AlphaFold2 (0.76). For the 72 multidomain proteins containing two conformational states with the maximum structural difference, the average TM-score of M-SADA is 0.87, which is 16.0% higher than that of AlphaFold2 (0.75). The head-to head comparison between the average TM-score is shown in **Figure 2 (a)**. For the 65 out of 72 multidomain proteins, the average TM-score of M-SADA is better than that of AlphaFold2. **Table 1** summarizes the number of multidomain proteins for which the M-SADA and AlphaFold2 models satisfy different TM-score cut-offs on two different conformational states. For 29 out of 72 proteins, the TM-scores of M-SADA models exceed 0.90 in both conformational states. However, the number of multidomain proteins with TM-scores above 0.90 in both conformational states for the AlphaFold2 models is 2.

**Table 1.** Number of multidomain proteins for which M-SADA and AlphaFold2 models satisfy the different TM-score cut-offs on two different conformational states.

| Method | The number of multidomain proteins |  |  |  |
| --- | --- | --- | --- | --- |
|  | TM-score<sub>s1, s2</sub> > 0.75 | TM-score<sub>s1, s2</sub> > 0.80 | TM-score<sub>s1, s2</sub> > 0.85 | TM-score<sub>s1, s2</sub> > 0.90 |
| M-SADA | 46 | 44 | 38 | 29 |
| AlphaFold2 | 14 | 7 | 4 | 2 |

*Note:* TM-score<sub>s1, s2</sub> > cut-off represents the TM-score of the best model on state 1 and the TM-score of the best model on state 2 are both greater than cut-off.

**Figure 2.**
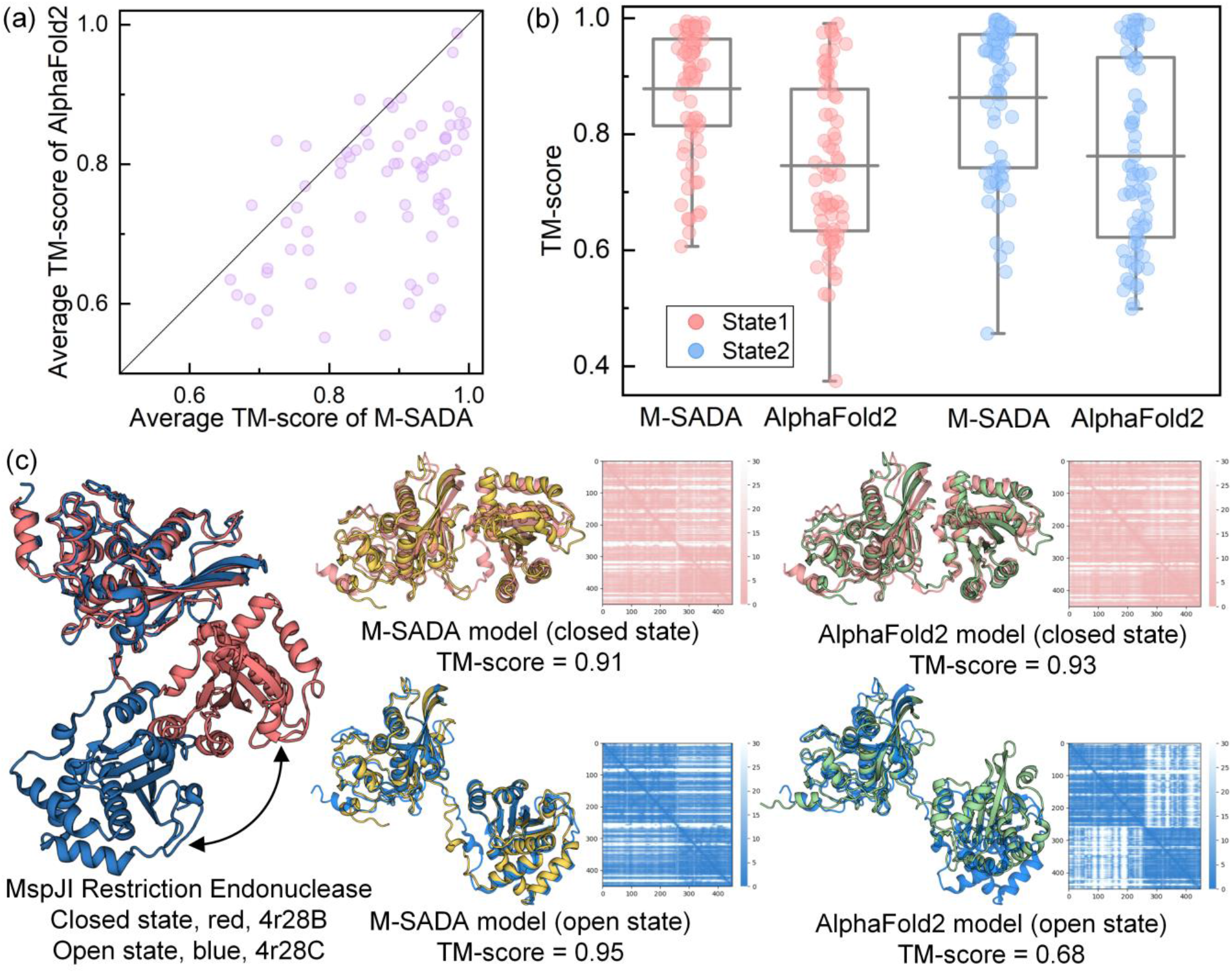
Comparison of M-SADA with AlphaFold2 on 72 multidomain proteins containing two conformational states with the maximum structural difference. (a) Head-to-head comparison between the average TM-scores of each multidomain protein generated by AlphaFold2 and M-SADA. (b) TM-score boxplot and distribution for different conformational states. The grey horizontal line in the box represents the average TM-scores. (c) A representative example is showing M-SADA performance in modelling different conformational states. The red and blue cartoons are native structures of different conformational states. The yellow and green cartoons represent the M-SADA model and AlphaFold2 model, respectively.

For a more intuitive analysis of the modelling performance on two representative conformational states, the TM-score boxplot and distribution for the two different states of each protein are shown in **Figure 2 (b)**. From this figure, we can draw the conclusion that M-SADA can accurately model two different conformational states on more proteins than state-of-the-art modelling method AlphaFold2. The result also validates that the multiple population-based evolutionary algorithm proposed in M-SADA is able to generate models with diversity, which enables M-SADA to explore multiple conformational states of multidomain proteins.

A representative example is shown in **Figure 2 (c)**, which is the MspJI restriction endonuclease in complex and belongs to a family of restriction enzymes that cleave DNA containing 5-methylcytosine (5mC) or 5-hydroxymethylcytosine (5hmC)^20^. In subunits A and B of the tetramer, the binding and cleavage domains are close together (‘closed’ conformation, protein 4r28B), and in subunits C and D they are farther apart with respect to each other (‘open’ conformation, protein 4r28C). For AlphaFold2, the ‘closed’ conformation is accurately modelled, whilst the subunits’ ‘open’ conformation is not modelled correctly. By contrast, M-SADA successfully modelled the ‘closed’ and ‘open’ conformations after the individual domain models were reassembled by M-SADA, achieving TM-scores of 0.91 and 0.95, respectively. These results indicate that M-SADA can probably be used to model different conformational states of multidomain proteins.

### Reassembly of multidomain protein from AlphaFold DB

Although AlphaFold2 has achieved remarkable success in modelling protein structures, the multidomain protein full-chain modelling accuracy appears to be worse on average than that for the constituent domains^1^. This shows the necessity of further effort on inter-domain orientation modelling for protein structure prediction. To test the effectiveness of M-SADA in modelling multidomain protein full-chain structures, M-SADA is used to reassemble 296 multidomain proteins with single conformational state, where the structures of individual domain models are from the AlphaFold2 full-chain model and 100% sequence identity cut-off is used to exclude templates for M-SADA. Here, we use the AlphaFold2 model to refer to the model deposited in AlphaFold DB, because the models deposited in AlphaFold DB are also predicted by AlphaFold2. **Table 2** summarizes the results of the full-chain models predicted by AlphaFold2 and reassembled by M-SADA and SADA. M-SADA-top1 represents the first model ranked by our developed DeepUMQA2^19^, which is currently one of state-of-the-art methods for protein monomer and complex model quality estimation. It ranks first in the protein complex interface residue accuracy estimation of CASP15. M-SADA-best represents the best model amongst the output models.

**Table 2.** Summary of modelling results for M-SADA, SADA and AlphaFold2 on 296 multidomain proteins.

| Method | Average TM-score |  |  |  |
| --- | --- | --- | --- | --- |
|  | pLDDT < 80 | 80 ≤ pLDDT < 90 | 90 ≤ pLDDT | All |
| M-SADA-best | 0.674 | 0.863 | 0.964 | 0.913 |
| M-SADA-top1 | 0.628 | 0.850 | 0.956 | 0.901 |
| SADA | 0.564 | 0.815 | 0.945 | 0.879 |
| AlphaFold2 | 0.524 | 0.801 | 0.940 | 0.868 |

*Note:* pLDDT < 80 represents the AlphaFold2 models with an average pLDDT <80.

Overall, the best models and top1 models of M-SADA obtain a higher TM-score than the models of AlphaFold2 and SADA. On average, the top1 M-SADA models achieve an average TM-score of 0.901, which is 3.8% higher than that of the AlphaFold2 models (0.868) and 2.5% higher than that of the SADA models (0.879). The average TM-score for the best models of M-SADA is 5.1% higher than that of the AlphaFold2 models. For the AlphaFold2 models with an average pLDDT ≥90, the average TM-score of the full-chain models assembled by M-SADA is slightly higher than that of AlphaFold2 models. However, for the AlphaFold2 models with an average pLDDT ≤80, the average TM-score of the top1 models of M-SADA achieves 0.628, which is 19.8% higher than that of AlphaFold2 (0.524) and 11.3% higher than that of SADA (0.564). The TM-score boxplot and distribution are illustrated in **Figure 3 (a)**. The head-to-head comparison between the TM-score of the top1 full-chain models of M-SADA and that of the AlphaFold2 and SADA models is shown in **Figure 3 (b)**. The results show that the quality of the AlphaFold2 models can be improved by M-SADA. In particular, for the 84 proteins with a TM-score of AlphaFold2 model <0.80, the average TM-score of the top1 models assembled by M-SADA is 0.729, which is 18.0% higher than that of AlphaFold2 (0.618). For the best models of M-SADA, they achieve a higher TM-score.

**Figure 3.**
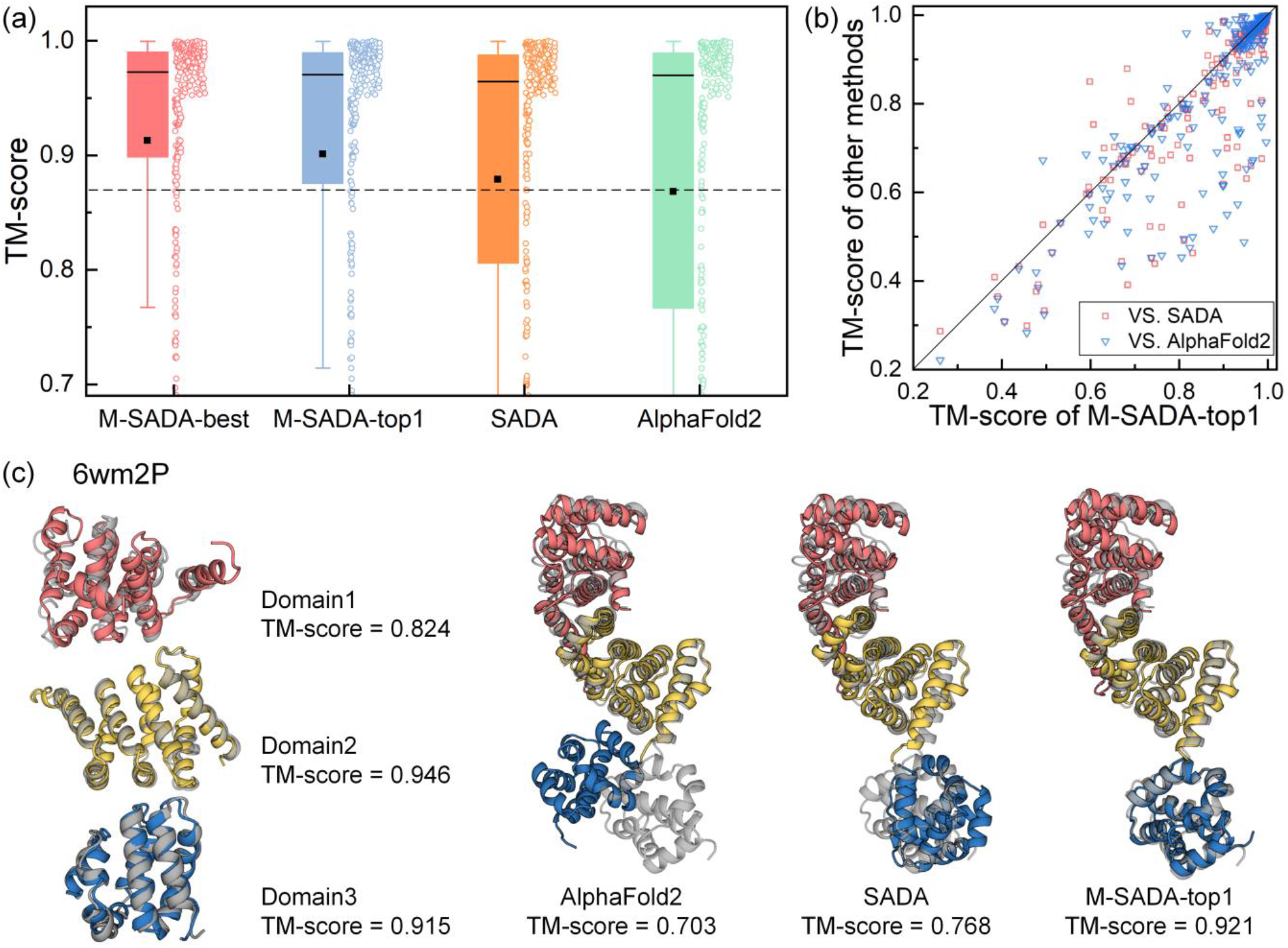
Comparison of the full-chain models assembled by M-SADA with that predicted by AlphaFold2. (a) TM-score boxplot and distribution for different methods. The black square and black horizontal line in the box represent the average and median TM-scores. (b) Head-to-head comparison between the TM-score of the top1 full-chain models of M-SADA and that of models predicted by AlphaFold2 and assembled by SADA. (c) A representative example is showing M-SADA can build better full-chain models. The grey cartoon represents the native structure, and different domains in the models are represented by different colours.

A representative example is shown in **Figure 3 (c)**. For 6wm2P with 3 domains, although AlphaFold2 accurately predicts all the domain models (TM-score = 0.824, 0.946 and 0.915), the TM-score of the full-chain model for AlphaFold2 is lower than the average TM-score of the constituent domains because the orientation of the third domain is not modelled correctly. In SADA, the orientation of the third domain is also not constructed correctly, resulting in a TM-score of 0.768 for the full-chain model. However, all domain orientations are modelled correctly by M-SADA, resulting in a TM-score of 0.921 for the top1 full-chain model.

These results also indicate that the accuracy of full-chain modelling is lower than the average accuracy of the constituent domains for some multidomain proteins, and M-SADA can probably improve the quality of AlphaFold2 models to a certain extent. Compared with SADA, M-SADA uses different types of templates and a more advanced inter-domain distance prediction network to generate multiple energy functions. Then, a multiple population-based evolutionary algorithm is used to generate more potential solutions. Therefore, the full-chain model assembled by M-SADA has a higher TM-score. To help determine in which scenarios M-SADA can be used to improve the quality of AlphaFold2 models, the relationship between the average pLDDT cut-off of the AlphaFold2 full-chain model and the proportion of the cases improved after M-SADA reassembly is shown in **Supplementary Figure S1**. Amongst the 296 multidomain proteins, for the AlphaFold2 models with an average pLDDT ≤90, more than 75% of the top1 models reassembled by M-SADA have higher TM-scores than before.

### Overall results of domain assembly on experimental domains

The accuracy of the single-domain model affects the performance of the domain assembly methods. In order to objectively compare the domain assembly performance of M-SADA with other methods, M-SADA is tested on 356 multidomain proteins and compared with three well-known domain assembly methods, namely, AIDA^11^, DEMO^12^ and SADA^14^.

We reassemble the individual domain structures partitioned from the experimental structure according to the domain boundary. The initial domain structure is randomly rotated and translated before assembly, and templates with a sequence identity >30% to the query are excluded. **Table 3** summarizes the modelling results of M-SADA and other methods.

**Table 3.**
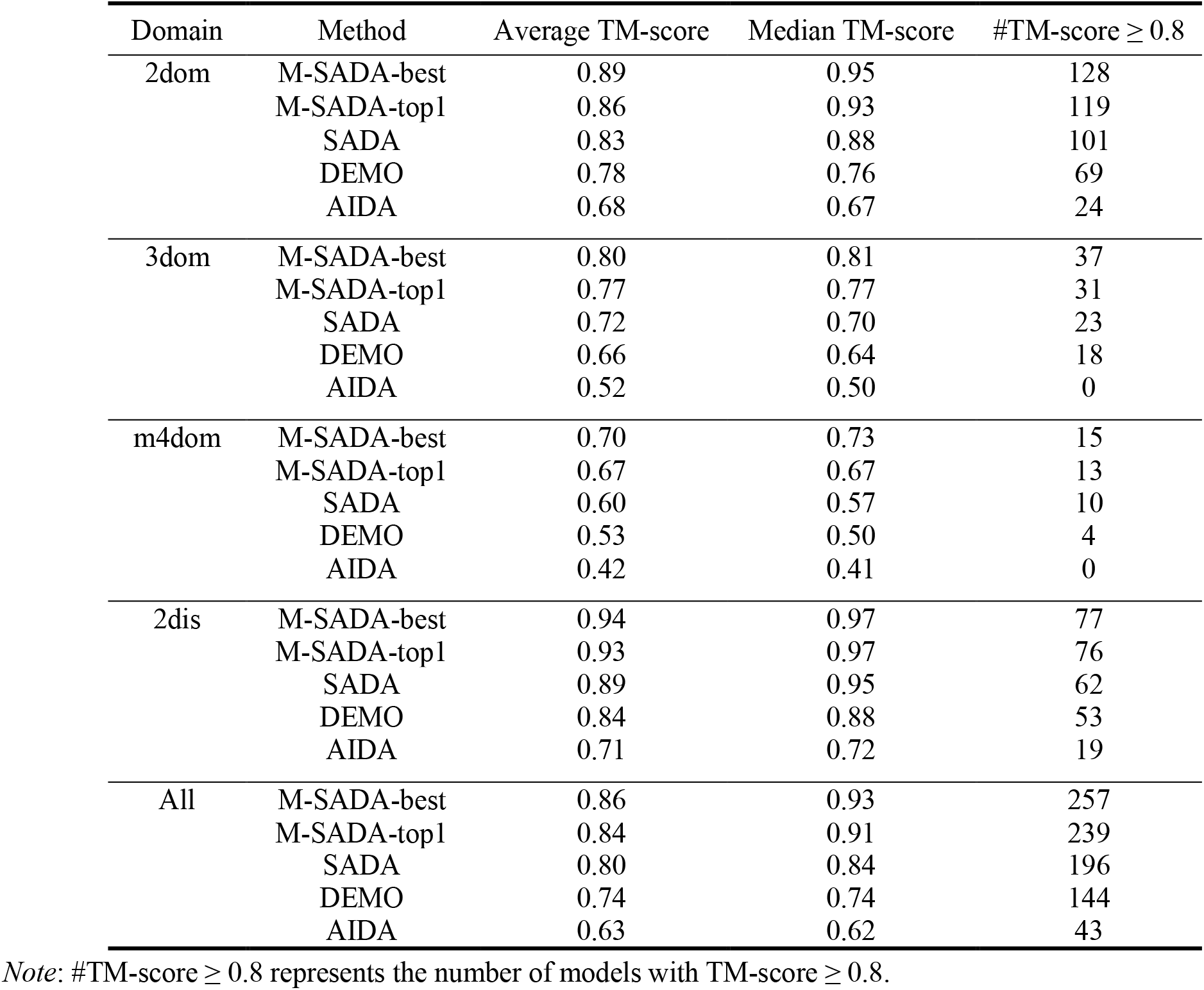
Summary of domain structure assembly by using experimentally solved domains on 356 test proteins.

For the top1 models of M-SADA, M-SADA obtains an average TM-score of 0.86 for 2dom, 0.77 for 3dom, 0.67 for m4dom and 0.93 for 2dis proteins. Overall, the average TM-score of M-SADA-top1 models is 0.84, which is 5.0% higher than that of the previously developed SADA (with *P*-value = 4.44e-11) and 13.5% higher than that of DEMO (with *P*-value = 3.17e-35). M-SADA accurately assembles (i.e. TM-score ≥0.8) 239 out of the 356 targets, accounting for 67.1% of the total, which is 12.0% and 26.7% more than that of SADA (55.1%) and DEMO (40.4%), respectively. In particular, for the m4dom proteins, the improvement in accuracy of the full-chain models is more significant, and the average TM-score of M-SADA is 11.7% higher than that of SADA. In addition, the results also indicate that the performance of M-SADA decreases when the number of domains increases, probably because the quality of the available templates decreases and the accuracy of the inter-residue distances predicted by DeepIDDP decreases when the number of domains increases, which affects the precision of the energy functions.

Nevertheless, the overall quality of the multidomain models is acceptable, and the *P*-values of SADA, DEMO and AIDA indicate statistically significant differences between the methods. The reason for such discrepancy is that M-SADA uses different types of templates and a more advanced inter-domain distance map to generate multiple energy functions that contain more potential orientations between domains.

### Evaluation of inter-domain distance prediction

In this section, we compare the performance of DeepIDDP in M-SADA with that of the distance prediction network GeomNet^21^ employed by SADA on 356 multidomain proteins. Here, all inter-domain residue pairs with a true distance of less than 15Å is considered because the number of contacts (true distance is less than 8Å) of inter-domain residual pair is small^22^. The comparison results are summarized in **Table 4**.

**Table 4.**
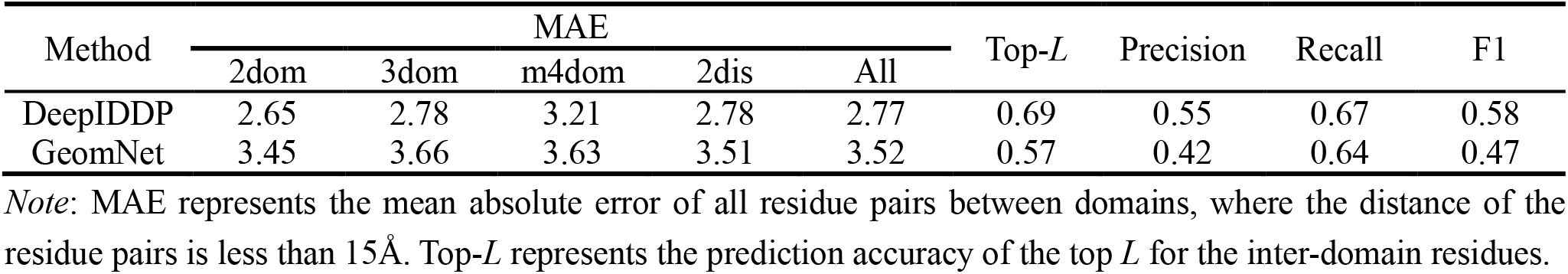
Summary of the results of inter-domain distance prediction for 356 multi-domain proteins.

Overall, DeepIDDP outperforms GeomNet on the multidomain protein test set. The mean absolute error (MAE) of DeepIDDP is 2.77Å, which is 21.3% lower than that of GeomNet (3.52Å). The top-*L* accuracy of DeepIDDP is 0.69, which is 21.0% higher than that of GeomNet (0.57). **Figure 4** shows the head-to-head comparison between the top-*L* accuracy of DeepIDDP and that of GeomNet on each test protein. In addition, we use three metrics, namely, precision, recall and F1, to comprehensively measure the predictive performance of the two distance predictors. The results listed in **Table 4** show that DeepIDDP outperforms GeomNet in inter-domain distance prediction.

**Figure 4.**
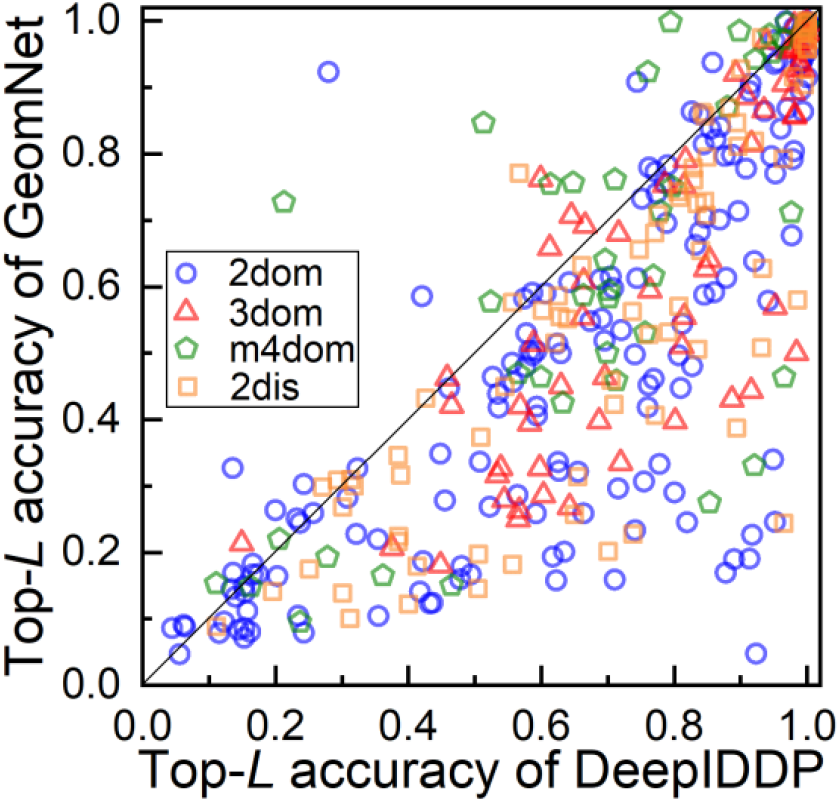
Head-to-head top-*L* accuracy comparison between DeepIDDP in M-SADA and GeomNet in SADA.

### Effect of homologous and analogous templates on M-SADA

Homologous and analogous templates are used to guide domain assembly in M-SADA. To further study the effects of analogues and homologues on the accuracy of domain assembly, M-SADA is used to assemble the full-chain structure of multidomain proteins according to the different templates. Here, M-SADA without using homologous templates is named M-SADA-w/o-H, and the M-SADA without using analogous templates is named M-SADA-w/o-A. In order to objectively study the effect of different templates on M-SADA, we only discuss the best models here.

For 296 out of the 356 test proteins, we discuss the effect of different templates on M-SADA, where other 60 proteins have no available homologous templates at a sequence identity cut-off of 30%. The domain assembly results of M-SADA, M-SADA-w/o-A and M-SADA-w/o-H on the 296 test proteins are summarized in **Supplementary Table S3**. Overall, the average TM-score of M-SADA is 0.87, which is 11.5% higher than that of M-SADA-w/o-A (0.78) and 10.1% higher than that of M-SADA-w/o-H (0.79), respectively. For an intuitive comparison of the effect, the TM-score boxplot and distribution in separate categories is depicted in **Figure 5(a)**, and the head-to-head comparison between the TM-scores of the models assembled by M-SADA that uses only analogous (or homologous) templates is shown in **Figure 5(b)**.

**Figure 5.**
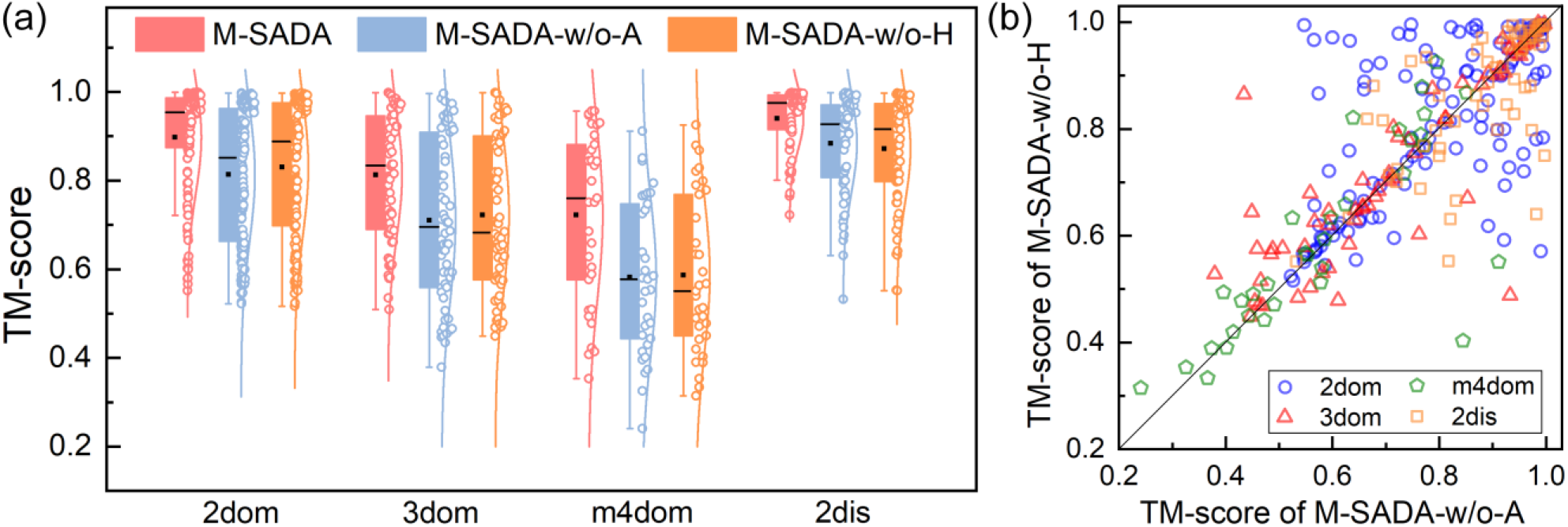
(a) Boxplot and distribution for the TM-score of the best models assembled by M-SADA, M-SADA-w/o-A and M-SADA-w/o-H on 296 test proteins. The black square and black horizontal line in the box represent the average and median TM-scores. (b) Head-to-head comparison between the TM-scores of the models assembled by M-SADA-w/o-A and M-SADA-w/o-H.

From **Figure 5(a)**, the average TM-score of M-SADA-w/o-H is slightly lower than that of M-SADA-w/o-A on the 2dis proteins, but slightly higher than that of M-SADA-w/o-A on the 2dom, 3dom and m4dom proteins. However, M-SADA achieves better TM-score than M-SADA-w/o-A and M-SADA-w/o-H. In particular, for 3dom and m4dom proteins, the average TM-scores for M-SADA are 0.81 and 0.72, respectively, which are 14.1% (0.71) and 24.1% (0.58) higher than those for M-SADA-w/o-A and 12.5% (0.72) and 22.0% (0.59) higher than those for M-SADA-w/o-H, respectively. **Figure 5(b)** shows that different types of templates may be complementary, and they can provide M-SADA with different types of template information. Thus, the combination of analogues and homologues can improve the modelling accuracy of multidomain proteins.

## Conclusion and discussion

The protein structure prediction problem has achieved remarkable progress through deep learning end-to-end techniques, in which high accurate models can be built for nearly all single-domain proteins and some multidomain proteins. However, compared with single-domain protein prediction, the full-chain modelling of multidomain proteins remains relatively unreliable. The modelling of multidomain protein full-chain structures remains an important problem that is largely ignored by mainstream computational biology. In addition, the domains of multidomain proteins frequently interact with each other, and therefore, multidomain proteins usually have multiple conformational states. The modelling of multidomain proteins with multiple conformational states is also essential for further studying protein functions and interaction mechanisms. However, fewer computational approaches have been developed to address this problem.

This work develops a multiple conformational states domain assembly method, M-SADA, which enables the modelling of different conformational states of multidomain proteins. The method starts with the recognition of analogous full-chain templates by structurally aligning component domains with proteins from MPDB, and the available homologous templates are identified from MPDB, PDB and AlphaFold DB. Based on the different templates, specific energy functions combined with deep learning predicted restraints are designed to guide domain assembly. Finally, a multiple population-based evolutionary algorithm is proposed to explore domain orientation based on multiple energy functions.

M-SADA is tested on 72 multidomain proteins with multiple conformational states, where the individual domain models are from the first model of AlphaFold2. Consequently, M-SADA correctly generates full-chain models with different states on the majority of multidomain proteins and achieves an average TM-score of 0.87, which is 16.0% higher than that of AlphaFold2 (0.75). M-SADA exhibits superior performance in modelling multiple conformational states of multidomain proteins mainly because of the following reasons: (i) it generates multiple conformational landscapes, and (ii) it achieves cooperation and optimization between different landscapes through a multiple population-based evolutionary algorithm. M-SADA is also applied to reassemble 296 human multidomain proteins from AlphaFold DB, and most cases achieve better models. In particular, for the 84 proteins with a TM-score <0.8 for the AlphaFold2 model, the average TM-score of the top1 models reassembled by M-SADA is 0.729, which is 18.0% higher than that of AlphaFold2 (0.618). The results show that M-SADA can complement to end-to-end protein structure prediction methods (e.g., AlphaFold2) to generate alternative or better models. M-SADA is also tested over a comprehensive set of 356 proteins with varying levels of continuous and discontinuous domain structures. The results show that M-SADA significantly outperforms state-of-the-art domain assembly methods. Meanwhile, the results of ablation experiment indicate that sequential homologues and structural analogues are complementary.

Despite the successes documented herein, M-SADA could be further improved in several aspects. Multiple MSAs are not considered by DeepIDDP. Therefore, the inter-domain distance map predicted by DeepIDDP lacks diversity. If diverse inter-domain distance maps can be predicted, the ability to model multidomain proteins with multiple conformational states will be likely improved. Currently, domain structures are kept rigid during M-SADA simulations, which cannot appropriately account for binding-induced structural changes, and thus, introducing backbone flexibility to domain assembly simulation provides the potential for local domain structure refinement. In addition, a gap still exists between the best model of M-SADA and the top1 model assessed by DeepUMQA2. Therefore, further improving the performance of model quality assessment for multidomain proteins is also the next direction of our research.

## Methods

### Templates for domain assembly

Templates are important for modelling protein structures. Structural information from templates is considerably more reliable than that from other sources, particularly when the target protein and the template are highly homologous. However, homologous templates that can be used to model the full-chain structure of multidomain proteins are often unavailable because most of the multidomain proteins have only single domain structures solved in PDB^12^. In practice, the structural space of protein-protein interfaces is small (<1000 structurally distinct interfaces)^5^. Similarly, the structural space of domain-domain interfaces is also limited. Therefore, structural analogues are suitable for multidomain protein modelling in some cases, because domains often interact in a similar way in the quaternary orientations if the domains have similar tertiary structures^14^. The comprehensive use of inter-domain interaction information provided by sequential homologues and structural analogues is necessary for domain assembly. Here, homologous templates are detected based on the input full-chain sequence, and analogous templates of the full-chain are identified according to the structural similarity between domain models and the proteins in MPDB. Different templates are used to construct multiple different energy functions for domain assembly, because they are complementary for modelling multidomain proteins to a certain extent^14^.

### Multidomain protein structure database update

Domains often interact with each other to perform biological functions, and thus, some multidomain proteins in the PDB are identical in sequence but different in structure. These diverse structures with the same sequence may also be important. Thus, we add multidomain protein structures with identical sequences but structural differences to MPDB as follows: (i) clustering all proteins in PDB with 100% sequence identity and then calculating the TM-scores between the proteins in each cluster, (ii) selecting the clusters with at least one pair of proteins with TM-score ≤ 0.85 and (iii) selecting the two proteins with the largest structural differences in each selected cluster to MPDB.

### Analogous template identification

Structurally analogous proteins comprise an important class of templates for building the full-chain structure of multidomain proteins. Here, analogous templates for full-chain structures are identified from MPDB through a two-stage procedure based on TM-align^23^, as shown in **Figure 6**.

**Figure 6.**
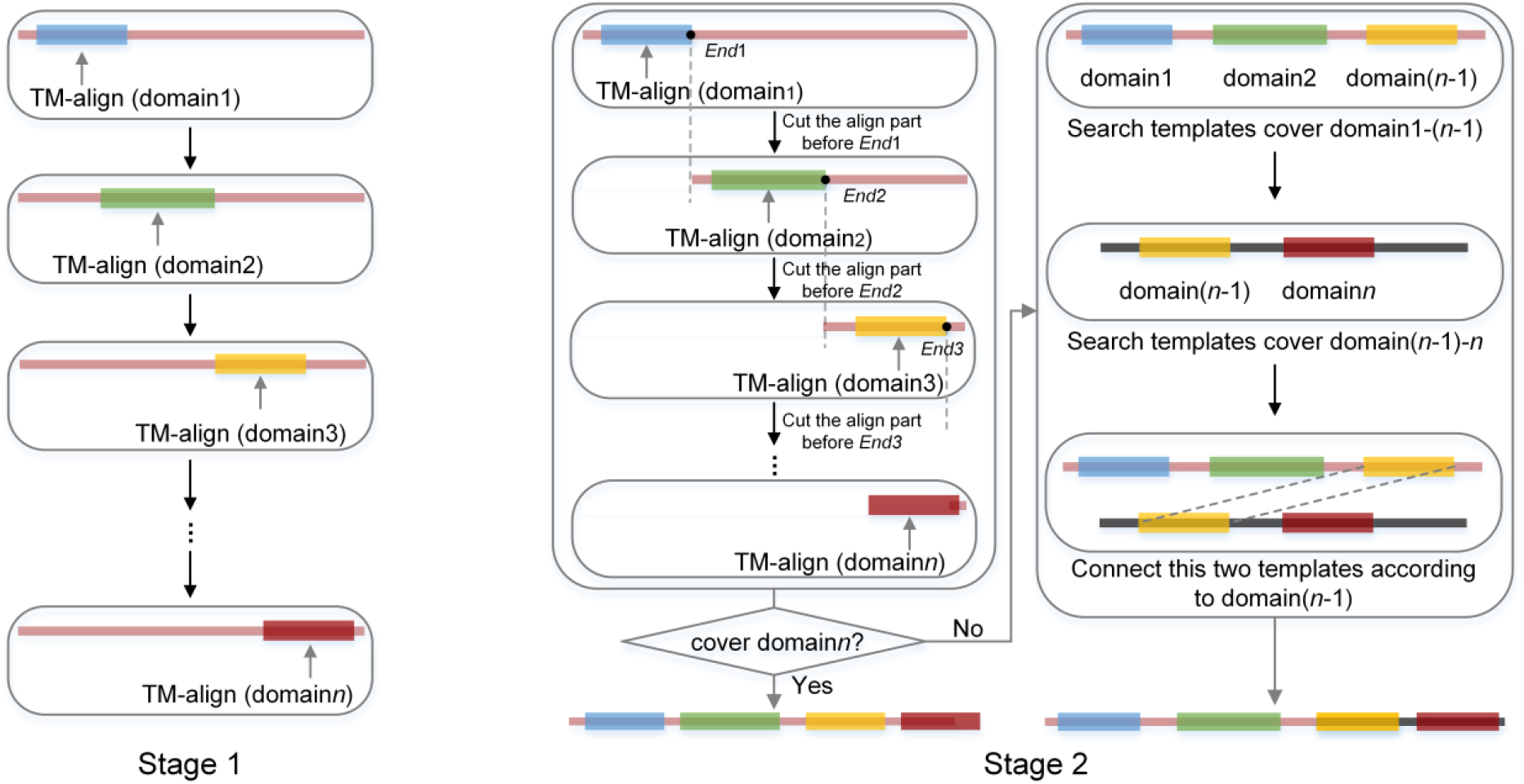
Process of analogous template identification.

In the first stage, individual domains of the query are structurally aligned with a template, regardless of the overlap between the alignments of different domains. A local similarity score *LS*_score_ is used to evaluate the quality of the template, and the top 200 templates with the highest *LS*_score_s are selected for the next stage, where *LS*_score_ is similarly defined as that in our previously developed SADA^14^.

In the second stage, the input individual domains are aligned on each template selected in the first stage using TM-align, where overlap is not allowed in the alignments of different domains to prevent structurally similar domains from being matched to the same part of the protein. The top five analogous templates with the highest *LS*_score_s in the second stage are selected for domain assembly. If the best template with the highest *LS*_score_ cannot cover all domains (e.g., one of the domains has the TM-score <0.5), the templates of the two broken parts will be independently detected. The templates that can cover domains1-(*n*-1) and the templates that can cover domains(*n*-1)-*n* are independently detected from the previously selected templates. The template with the highest *LS*_score_ for domains1- (*n*-1) and the template with the highest *LS*_score_ for domains(*n*-1)-*n* are selected for domain assembly.

### Homologous template recognition

Homologous proteins play a pivotal role in predicting the structure of proteins that have not been studied experimentally^24^, primarily because homologous proteins share a common ancestor, and thus, they usually have similar structures in homologous sequence regions. However, available protein structures deposited in the PDB are limited (about 200,000 publicly available structures as of January 2023)^25^, accounting for less than 0.1% of the protein sequence database UniProtKB/TrEMBL^26^, resulting in some homologous proteins not being available for structural modelling. Recently, DeepMind and EMBL-European Bioinformatics Institute collaborates to create a new data resource, AlphaFold DB, which greatly expands the structural coverage of the known protein sequence space^6^. This makes it possible for AlphaFold DB to complement PDB with respect to the modelling full-chain structure of multidomain protein. Therefore, we introduce AlphaFold DB to compensate for the lack of homologous structures available in PDB. The process of detecting the homologous proteins of the target sequence can be divided into four stages that correspond to the searching for four structure databases (MPDB, PDB, AlphaFold DB90 and AlphaFold DB) through HMMER program. The details can be found in **Supplementary Text S1**.

### Inter-domain distance prediction neural network

To our knowledge, few distance prediction methods are optimized specifically for the distance of the inter-domain residue pairs of multidomain proteins. Currently, most multidomain protein structure assembly methods use inter-domain distances extracted directly from the distances predicted by methods developed for common distance prediction, possibly reducing ability to capture orientations between domains.

In this work, our recently developed inter-domain distance prediction network (DeepIDDP) is introduced into M-SADA. In DeepIDDP, a data enhancement strategy, called domain-pair multiple sequence alignment (DPMSA), is firstly proposed to enhance the inter-domain coevolutionary information of multidomain proteins. Then, two new features, namely, the inter-domain residue coupling score and the inter-domain average contact potential (ICOP), are designed to focus on inter-domain residue pairs. Finally, a deep network that combines the attention mechanism and a deep residual block is constructed to predict the inter-domain residue distance distribution of multidomain proteins. The details of DeepIDDP can be referenced in [18].

### Energy functions for domain assembly

For multidomain proteins, capturing the interactions between domains is difficult for a single template in some cases because domains often interact with each other to perform more complex functions, resulting in a diversity of orientations between domains. Here, we design multiple energy functions based on different template information to guide domain assembly.

For homologue- or analogue-based populations, we design different energy functions 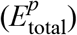 for each population based on different templates to guide domain assembly, where *p* represents the energy function used in the *p*-th population. 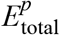 is defined as follows:

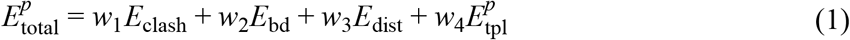

where each energy term is described in **Supplementary Test S2**.

For the hybrid population, we use all the detected template information to design a hybrid energy function 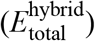 to guide domain assembly. 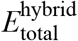 is defined as follows:

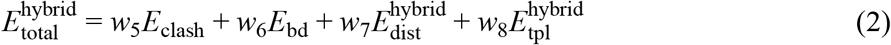

where each energy term is described in **Supplementary Test S2**.

### Multiple population-based evolutionary algorithm for domain assembly

For different energy functions, a multiple population-based evolutionary algorithm is proposed to explore optimal solutions, in which population diversity can be maintained because different populations can be located in different search spaces. Solutions amongst different populations can be exchanged rather than independently optimized, enabling the finding of more potential solutions efficiently. The flowchart of the multiple population-based evolutionary algorithm is shown in **Supplementary Figure S2** and described in **Supplementary Text S3**. The proposed algorithm can further improve domain assembly accuracy while ensuring the diversity of domain assembly results.

The same number of populations are set based on the number of energy functions. In each population, a two-stage differential evolution algorithm is proposed to explore and exploit the optimal solution under the guidance of the corresponding energy function. During the simulation, the solutions amongst different populations are exchanged for each learning period generation, where the best *k* solutions from each population are used to replace the worst *P* × *k* solutions in the hybrid population. Here, *P* is the number of populations in addition to the hybrid population. Then, the worst *k* solutions of each population are replaced with the best *k* solutions of the hybrid population. When the populations converge, the solution with the lowest 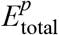 in each population is selected and ranked by DeepUMQA2 for output.

For the hybrid population, inspired by ultrafast shape recognition^27, 28^, three different solutions are selected based on the cosine similarity of the solutions because multiple template information is utilized in the hybrid population and different stable solutions may be able to increase the diversity of solutions. The rules for the selection of solutions in the hybrid population are as follows: (1) the solution with the lowest 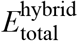 is selected, named *S*_hybrid, best_, (2) the solution with the minimum cosine similarity to *S*_hybrid, best_ is selected, name *S*_hybrid, Mbest_ and (3) the solution with the minimum cosine similarity to *S*_hybrid, Mbest_ is selected, named *S*_hybrid, MMbest_. When the cosine similarity amongst the solutions is smaller, the structural difference amongst the models generated by these solutions is greater.

The parameter setting of the multiple population-based evolutionary algorithm is described in **Supplementary Text S3**.

## Data availability

All data needed to evaluate the conclusions in the paper are present in the manuscript and Supplementary Information. The data are also available upon request from the corresponding author.

## Code availability

The online server of M-SADA is made freely available at http://zhanglab-bioinf.com/M-SADA/.

## Acknowledgments

This work is supported by the “New Generation Artificial Intelligence” major project of Science and Technology Innovation 2030 of the Ministry of Science and Technology of the People’s Republic of China (2022ZD0115100), the National Nature Science Foundation of China (62173304 and 62203389), and the Key Project of Zhejiang Provincial Natural Science Foundation of China (LZ20F030002). We thank Fengqi Ge for his discussions and feedbacks.

## Author contributions

G.Z. conceived and designed research. C.P., X.Z. and M.H. wrote algorithm and performed the experiments. G.Z. and C.P. developed DeepIDDP program. G.Z. and J.L. designed and developed DeepUMQA2 program. G.Z. and C.P. analyzed data and developed the server. S.L. helped supervise the research. G.Z. and C.P. wrote the manuscript, and all authors proofread the manuscript.

