## Supplementary information for "Multiple conformational states assembly of multidomain proteins using evolutionary algorithm based on structural analogues and sequential homologues"

### Supplementary Figures

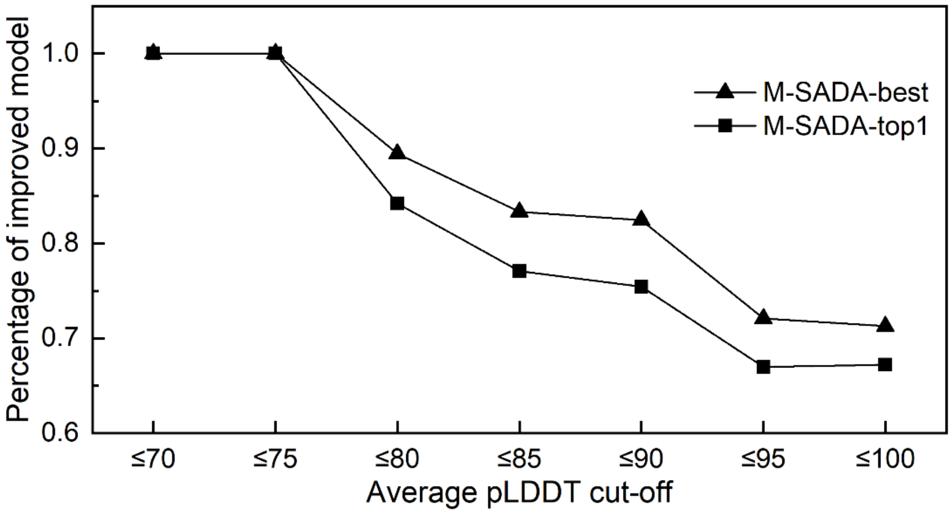


**Figure S1.** The relationship between different average pLDDT cut-offs of the AlphaFold2 full-chain model and proportions of the cases improved after M-SADA reassembly. AlphaFold produces a per-residue estimate of its confidence on a scale from 0~100. This confidence measure is called pLDDT. Average pLDDT represents a full-chain model estimate by calculating the average of pLDDT for all residues. M-SADA-best represents the model with the highest TM-score. M-SADA-top1 represents the ranked one model, which is ranked by DeepUMQA2.

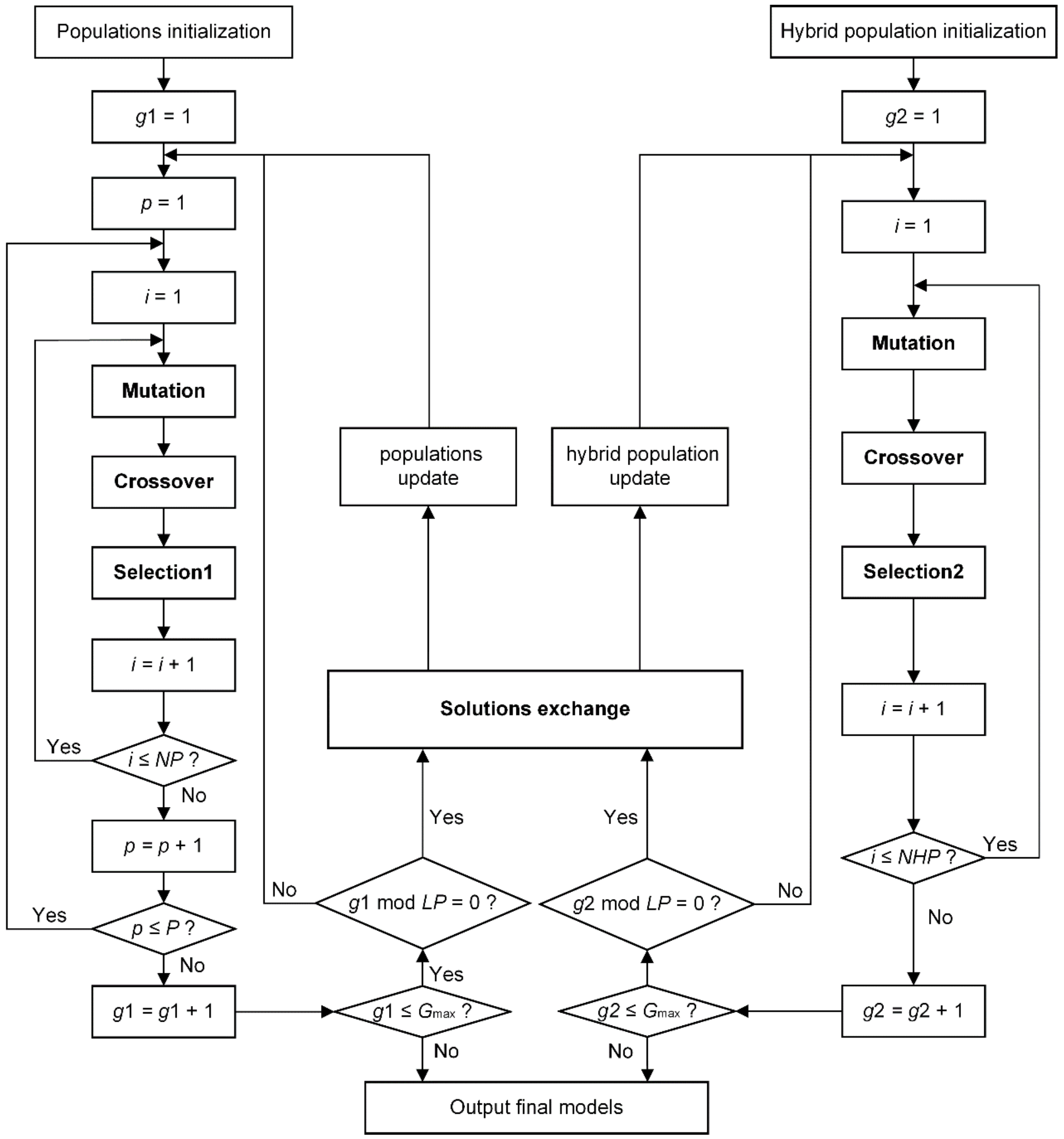


**Figure S2.** The flowchart of the multiple population-based evolutionary algorithm. First, each population is initialized. In the populations initialization and hybrid population initialization, the initial solutions are generated randomly. After crossover and mutation operations, new solutions are generated, and then a selection operation is performed to determine whether the new solutions can replace the old solutions in the populations (including hybrid population). The crossover, mutation and selection operations are iterated in each population (including hybrid population). The potential solutions in each population (including hybrid population) are exchanged to achieve interaction between populations and hybrid population when the number of iterations satisfies the learning period (*LP*). The above process is performed iteratively, and the final solutions are output when the number of iterations is satisfied.

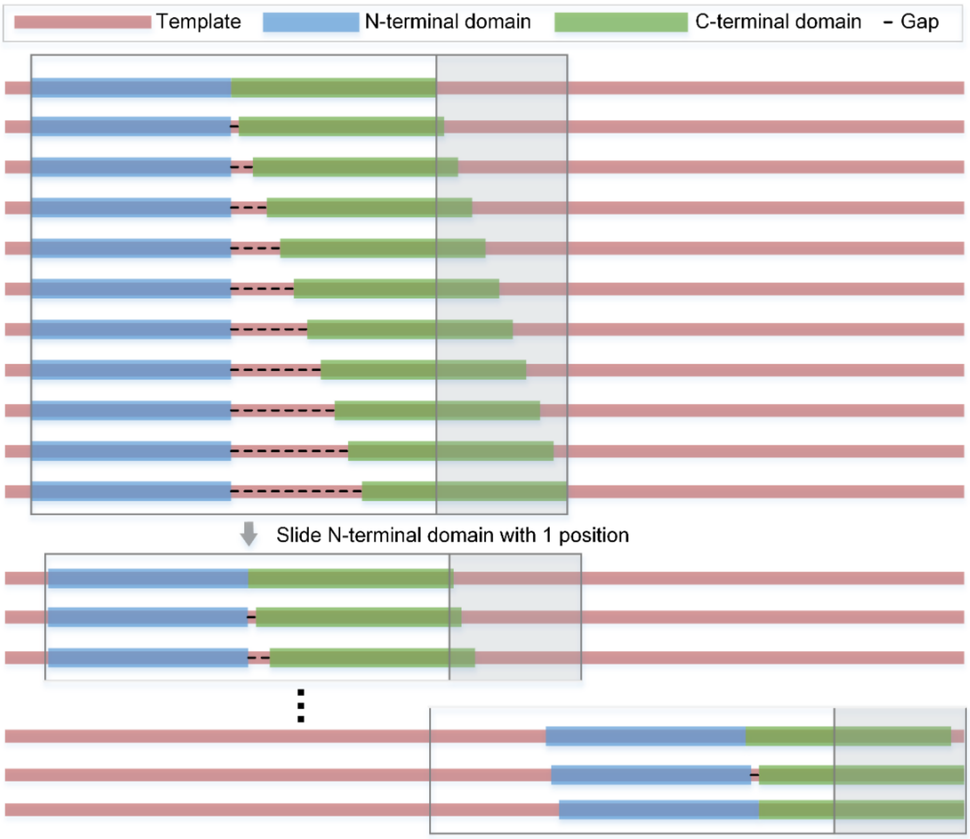


**Figure S3.** Sliding-window procedure for domain-template alignment and template-based domain positions build. In this procedure, the N-terminal domain is superposed with every position along the template, where at each position, the C-terminal domain is allowed to superpose in the remaining regions of the template at a maximum of 10 residues away from the N domain. The alignment with the highest average TM-score is finally selected to build the template-based domain positions.

### Supplementary Tables

**Table S1.** Summary of 72 multidomain proteins with multiple conformational states. State1 and state2 represent the two highly distinct conformations in each multidomain protein. State1&2 represents the average TM-score of the two highly distinct conformations. All represents the average TM-score of all conformations.

| Methods | Average TM-score | | | |
|---|---|---|---|---|
| | State1 | State2 | State1&2 | All |
| M-SADA | 0.88 | 0.86 | 0.87 | 0.87 |
| AlphaFold2 | 0.75 | 0.76 | 0.75 | 0.77 |

**Table S2.** Details of 72 multidomain proteins with multiple conformational states. The details of each multidomain protein cluster are shown in **Table S2.1** to **Table S2.72**. The bolded bold indicates the two most structurally different conformations.

**Table S2.1** Multidomain protein cluster 1.

| PDB ID | Best TM-score | |
|---|---|---|
| | M-SADA | AlpahFold2 |
| **2wg5F** | 0.989 | 0.983 |
| 2wg5G | 0.943 | 0.643 |
| 2wg5I | 0.943 | 0.643 |
| 2wg5J | 0.989 | 0.982 |
| **2wg5K** | 0.942 | 0.643 |
| 2wg5L | 0.989 | 0.982 |
| Average TM-score | 0.966 | 0.812 |

**Table S2.2** Multidomain protein cluster 2.

| PDB ID | Best TM-score | |
|---|---|---|
| | M-SADA | AlpahFold2 |
| **4uc4A** | 0.815 | 0.944 |
| **4uc4B** | 0.967 | 0.820 |
| Average TM-score | 0.891 | 0.882 |

**Table S2.3** Multidomain protein cluster 3.

| PDB ID | Best TM-score | |
|---|---|---|
| | M-SADA | AlpahFold2 |
| **3on0B** | 0.936 | 0.619 |
| **3on0C** | 0.962 | 0.654 |
| Average TM-score | 0.949 | 0.637 |

**Table S2.4** Multidomain protein cluster 4.

| PDB ID | Best TM-score | |
|---|---|---|
| | M-SADA | AlpahFold2 |
| **1m5q1** | 0.972 | 0.872 |
| 1m5q2 | 0.973 | 0.874 |
| 1m5qB | 0.954 | 0.604 |
| 1m5qC | 0.956 | 0.600 |
| 1m5qD | 0.955 | 0.607 |
| 1m5qE | 0.954 | 0.606 |
| 1m5qF | 0.956 | 0.599 |
| 1m5qG | 0.957 | 0.605 |
| 1m5qL | 0.973 | 0.865 |
| 1m5qO | 0.956 | 0.600 |
| 1m5qP | 0.954 | 0.609 |
| 1m5qQ | 0.957 | 0.605 |
| 1m5qR | 0.953 | 0.598 |
| 1m5qS | 0.955 | 0.600 |
| **1m5qT** | 0.955 | 0.598 |
| 1m5qU | 0.956 | 0.602 |
| Average TM-score | 0.958 | 0.653 |

**Table S2.5** Multidomain protein cluster 5.

| PDB ID | Best TM-score | |
|---|---|---|
| | M-SADA | AlpahFold2 |
| 4gkjM | 0.896 | 0.911 |
| **4gkkM** | 0.890 | 0.903 |
| **4v4gGM** | 0.744 | 0.701 |
| Average TM-score | 0.843 | 0.838 |

**Table S2.6** Multidomain protein cluster 6.

| PDB ID | Best TM-score | |
|---|---|---|
| | M-SADA | AlpahFold2 |
| 2hzaA | 0.913 | 0.672 |
| **2hzvA** | 0.857 | 0.956 |
| 2hzvB | 0.934 | 0.649 |
| 2hzvC | 0.875 | 0.962 |
| 2hzvD | 0.916 | 0.649 |
| 2hzvE | 0.859 | 0.952 |
| 2hzvF | 0.917 | 0.647 |
| 2hzvG | 0.840 | 0.959 |
| **2hzvH** | 0.940 | 0.646 |
| Average TM-score | 0.895 | 0.788 |

**Table S2.7** Multidomain protein cluster 7.

| PDB ID | Best TM-score | |
|---|---|---|
| | M-SADA | AlpahFold2 |
| **3gn5A.pdb** | 0.965 | 0.895 |
| 3o9xA.pdb | 0.967 | 0.778 |
| **3o9xB.pdb** | 0.966 | 0.779 |
| Average TM-score | 0.966 | 0.818 |

**Table S2.8** Multidomain protein cluster 8.

| PDB ID | Best TM-score | |
|---|---|---|
| | M-SADA | AlpahFold2 |
| **2bj3C** | 0.897 | 0.615 |
| 2bj7B | 0.872 | 0.967 |
| 2bj8B | 0.872 | 0.966 |
| **2bj9B** | 0.869 | 0.965 |
| Average TM-score | 0.877 | 0.878 |

**Table S2.9** Multidomain protein cluster 9.

| PDB ID | Best TM-score | |
|---|---|---|
| | M-SADA | AlpahFold2 |
| 1iq5A.pdb | 0.665 | 0.711 |
| 1lvcD.pdb | 0.553 | 0.516 |
| 1lvcE.pdb | 0.542 | 0.509 |
| 1lvcF.pdb | 0.545 | 0.515 |
| 1wrzA.pdb | 0.872 | 0.803 |
| 2bkiB.pdb | 0.838 | 0.694 |
| 2r28A.pdb | 0.907 | 0.654 |
| 2r28B.pdb | 0.892 | 0.669 |
| **2w73A.pdb** | 0.916 | 0.679 |
| 2w73B.pdb | 0.909 | 0.705 |
| 2w73E.pdb | 0.870 | 0.671 |
| 3oxqA.pdb | 0.748 | 0.691 |
| 3oxqC.pdb | 0.748 | 0.698 |
| 4l79B.pdb | 0.472 | 0.470 |
| 4ovnA.pdb | 0.452 | 0.537 |
| 4ovnB.pdb | 0.467 | 0.550 |
| **4ovnC.pdb** | 0.456 | 0.535 |
| 4ovnD.pdb | 0.463 | 0.539 |
| 4ovnE.pdb | 0.461 | 0.525 |
| 4q5uA.pdb | 0.790 | 0.754 |
| 4umoC.pdb | 0.564 | 0.561 |
| 4umoD.pdb | 0.561 | 0.557 |
| 4v0cC.pdb | 0.557 | 0.554 |
| 4v0cD.pdb | 0.555 | 0.551 |
| 5v03R.pdb | 0.550 | 0.561 |
| 6b8lB.pdb | 0.566 | 0.554 |
| 6b8lD.pdb | 0.568 | 0.560 |
| 6b8lF.pdb | 0.570 | 0.556 |
| 6b8mB.pdb | 0.572 | 0.558 |
| 6b8mF.pdb | 0.574 | 0.565 |
| 6b8nB.pdb | 0.574 | 0.560 |
| 6b8nF.pdb | 0.573 | 0.564 |
| 6b8pB.pdb | 0.572 | 0.557 |
| 6b8pF.pdb | 0.572 | 0.561 |
| 6b8qD.pdb | 0.561 | 0.552 |
| 6b8qF.pdb | 0.567 | 0.555 |
| 6mbaB.pdb | 0.455 | 0.542 |
| 6n5wC.pdb | 0.561 | 0.540 |
| 7cr3C.pdb | 0.567 | 0.560 |
| 7cr3E.pdb | 0.567 | 0.560 |
| 7cr3F.pdb | 0.567 | 0.560 |
| 7cr3H.pdb | 0.567 | 0.560 |
| 7cr4C.pdb | 0.564 | 0.552 |
| 7cr4E.pdb | 0.564 | 0.552 |
| 7cr4G.pdb | 0.564 | 0.552 |
| 7cr4H.pdb | 0.564 | 0.552 |

| | | |
|---|---|---|
| 7cr7C.pdb | 0.570 | 0.556 |
| 7cr7E.pdb | 0.570 | 0.556 |
| 7cr7F.pdb | 0.570 | 0.556 |
| 7cr7H.pdb | 0.570 | 0.556 |
| 7kl5A.pdb | 0.609 | 0.672 |
| Average TM-score | 0.609 | 0.584 |

**Table S2.10** Multidomain protein cluster 10.

| PDB ID | Best TM-score | |
|---|---|---|
| | M-SADA | AlpahFold2 |
| **5dmjA** | 0.770 | 0.640 |
| **5ihlF** | 0.777 | 0.617 |
| Average TM-score | 0.774 | 0.629 |

**Table S2.11** Multidomain protein cluster 11.

| PDB ID | Best TM-score | |
|---|---|---|
| | M-SADA | AlpahFold2 |
| **1vf5D** | 0.803 | 0.775 |
| **2d2cQ** | 0.856 | 0.846 |
| Average TM-score | 0.829 | 0.810 |

**Table S2.12** Multidomain protein cluster 12.

| PDB ID | Best TM-score | |
|---|---|---|
| | M-SADA | AlpahFold2 |
| **4tu7A** | 0.950 | 0.670 |
| **4tu7B** | 0.902 | 0.568 |
| Average TM-score | 0.926 | 0.619 |

**Table S2.13** Multidomain protein cluster 13.

| PDB ID | Best TM-score | |
|---|---|---|
| | M-SADA | AlpahFold2 |
| **1qr4A** | 0.947 | 0.866 |
| **1qr4B** | 0.740 | 0.919 |
| Average TM-score | 0.844 | 0.893 |

**Table S2.14** Multidomain protein cluster 14.

| PDB ID | Best TM-score | |
|---|---|---|
| | M-SADA | AlpahFold2 |
| **4a0zA** | 0.918 | 0.925 |
| 4a12A | 0.613 | 0.613 |
| **4a12D** | 0.612 | 0.612 |
| Average TM-score | 0.714 | 0.717 |

**Table S2.15** Multidomain protein cluster 15.

| PDB ID | Best TM-score | |
|---|---|---|
| | M-SADA | AlpahFold2 |
| **1ek8A** | 0.731 | 0.637 |
| 1zn0A | 0.734 | 0.694 |
| 1zn1A | 0.796 | 0.839 |
| **2rdo8** | 0.930 | 0.609 |
| Average TM-score | 0.798 | 0.694 |

**Table S2.16** Multidomain protein cluster 16.

| PDB ID | Best TM-score | |
|---|---|---|
| | M-SADA | AlpahFold2 |
| **1ikuA** | 0.607 | 0.565 |
| **1jsaA** | 0.870 | 0.867 |
| Average TM-score | 0.739 | 0.716 |

**Table S2.17** Multidomain protein cluster 17.

| PDB ID | Best TM-score | |
|---|---|---|
| | M-SADA | AlpahFold2 |
| **2jheA** | 0.920 | 0.674 |
| 2jheB | 0.826 | 0.844 |
| **2jheD** | 0.944 | 0.913 |
| Average TM-score | 0.897 | 0.811 |

**Table S2.18** Multidomain protein cluster 18.

| PDB ID | Best TM-score | |
|---|---|---|
| | M-SADA | AlpahFold2 |
| **2fjrA** | 0.890 | 0.620 |
| **2fjrB** | 0.939 | 0.581 |
| Average TM-score | 0.914 | 0.600 |

**Table S2.19** Multidomain protein cluster 19.

| PDB ID | Best TM-score | |
|---|---|---|
| | M-SADA | AlpahFold2 |
| **2a38A** | 0.938 | 0.675 |
| 2a38B | 0.972 | 0.511 |
| **2a38C** | 0.980 | 0.508 |
| Average TM-score | 0.963 | 0.565 |

**Table S2.20** Multidomain protein cluster 20.

| PDB ID | Best TM-score | |
|---|---|---|
| | M-SADA | AlpahFold2 |
| **1bccE** | 0.959 | 0.779 |
| **2bccE** | 0.911 | 0.814 |
| 3bccE | 0.916 | 0.804 |
| Average TM-score | 0.928 | 0.799 |

**Table S2.21** Multidomain protein cluster 21.

| PDB ID | Best TM-score | |
|---|---|---|
| | M-SADA | AlpahFold2 |
| **4hyeA** | 0.957 | 0.752 |
| **4hyeB** | 0.830 | 0.850 |
| Average TM-score | 0.893 | 0.801 |

**Table S2.22** Multidomain protein cluster 22.

| PDB ID | Best TM-score | |
|---|---|---|
| | M-SADA | AlpahFold2 |
| **4n9hA** | 0.630 | 0.630 |
| 4n9hB | 0.754 | 0.674 |
| **4n9iA** | 0.686 | 0.639 |
| 4n9iB | 0.636 | 0.636 |
| 4n9iC | 0.686 | 0.640 |
| 4n9iD | 0.638 | 0.636 |
| 6b6hG | 0.957 | 0.970 |
| 6b6hH | 0.945 | 0.962 |
| Average TM-score | 0.742 | 0.723 |

**Table S2.23** Multidomain protein cluster 23.

| PDB ID | Best TM-score | |
|---|---|---|
| | M-SADA | AlpahFold2 |
| **3dkxC** | 0.985 | 0.864 |
| **4u87B** | 0.821 | 0.927 |
| 4u87C | 0.987 | 0.875 |
| Average TM-score | 0.931 | 0.888 |

**Table S2.24** Multidomain protein cluster 24.

| PDB ID | Best TM-score | |
|---|---|---|
| | M-SADA | AlpahFold2 |
| **3q4iA** | 0.949 | 0.945 |
| **3q4iB** | 0.727 | 0.696 |
| Average TM-score | 0.838 | 0.820 |

**Table S2.25** Multidomain protein cluster 25.

| PDB ID | Best TM-score | |
|---|---|---|
| | M-SADA | AlpahFold2 |
| **2h6cA** | 0.827 | 0.678 |
| **2h6cB** | 0.710 | 0.728 |
| Average TM-score | 0.768 | 0.703 |

**Table S2.26** Multidomain protein cluster 26.

| PDB ID | Best TM-score | |
|---|---|---|
| | M-SADA | AlpahFold2 |
| **4if4A** | 0.717 | 0.789 |
| 4if4B | 0.717 | 0.788 |
| 4if4C | 0.972 | 0.659 |
| **4if4D** | 0.972 | 0.660 |
| Average TM-score | 0.844 | 0.724 |

**Table S2.27** Multidomain protein cluster 27.

| PDB ID | Best TM-score | |
|---|---|---|
| | M-SADA | AlpahFold2 |
| 4qhkJ | 0.941 | 0.627 |
| 4qhkL | 0.947 | 0.632 |
| 4qhkN | 0.950 | 0.624 |
| **4qhkP** | 0.948 | 0.619 |
| 4qhlB | 0.927 | 0.613 |
| **4qhlD** | 0.956 | 0.544 |
| Average TM-score | 0.945 | 0.610 |

**Table S2.28** Multidomain protein cluster 28.

| PDB ID | Best TM-score | |
|---|---|---|
| | M-SADA | AlpahFold2 |
| **4bh9A** | 0.779 | 0.755 |
| **4bhpA** | 0.853 | 0.820 |
| Average TM-score | 0.816 | 0.787 |

**Table S2.29** Multidomain protein cluster 29.

| PDB ID | Best TM-score | |
|---|---|---|
| | M-SADA | AlpahFold2 |
| 3dppB | 0.971 | 0.856 |
| **4hybB** | 0.963 | 0.834 |
| 4jwdB | 0.971 | 0.852 |
| **7n6jA** | 0.978 | 0.932 |
| Average TM-score | 0.971 | 0.868 |

**Table S2.30** Multidomain protein cluster 30.

| PDB ID | Best TM-score | |
|---|---|---|
| | M-SADA | AlpahFold2 |
| **1jn6B** | 0.869 | 0.560 |
| **1jnhH** | 0.891 | 0.550 |
| Average TM-score | 0.880 | 0.555 |

**Table S2.31** Multidomain protein cluster 31.

| PDB ID | Best TM-score | |
|---|---|---|
| | M-SADA | AlpahFold2 |
| 4s04A.pdb | 0.984 | 0.975 |
| 4s04B.pdb | 0.888 | 0.556 |
| 4s04E.pdb | 0.963 | 0.967 |
| 4s04F.pdb | 0.952 | 0.550 |
| 4s05A.pdb | 0.981 | 0.965 |
| 4s05B.pdb | 0.837 | 0.557 |
| Average TM-score | 0.934 | 0.762 |

**Table S2.32** Multidomain protein cluster 32.

| PDB ID | Best TM-score | |
|---|---|---|
| | M-SADA | AlpahFold2 |
| **2nueA** | 0.898 | 0.727 |
| **4m2zA** | 0.934 | 0.960 |
| 4m30A | 0.937 | 0.964 |
| Average TM-score | 0.923 | 0.884 |

**Table S2.33** Multidomain protein cluster 33.

| PDB ID | Best TM-score | |
|---|---|---|
| | M-SADA | AlpahFold2 |
| **3u4mA** | 0.979 | 0.975 |
| **7a5fn** | 0.725 | 0.721 |
| 7a5gn | 0.732 | 0.728 |
| 7a5jn | 0.727 | 0.725 |
| Average TM-score | 0.791 | 0.787 |

**Table S2.34** Multidomain protein cluster 34.

| PDB ID | Best TM-score | |
|---|---|---|
| | M-SADA | AlpahFold2 |
| 6b6hA | 0.970 | 0.933 |
| **6govU** | 0.974 | 0.920 |
| **6pb5A** | 0.738 | 0.737 |
| Average TM-score | 0.894 | 0.863 |

**Table S2.35** Multidomain protein cluster 35.

| PDB ID | Best TM-score | |
|---|---|---|
| | M-SADA | AlpahFold2 |
| **3bdnA** | 0.913 | 0.822 |
| **3bdnB** | 0.913 | 0.627 |
| Average TM-score | 0.913 | 0.725 |

**Table S2.36** Multidomain protein cluster 36.

| PDB ID | Best TM-score | |
|---|---|---|
| | M-SADA | AlpahFold2 |
| **2q0oA** | 0.982 | 0.729 |
| **2q0oB** | 0.973 | 0.706 |
| Average TM-score | 0.977 | 0.718 |

**Table S2.37** Multidomain protein cluster 37.

| PDB ID | Best TM-score | |
|---|---|---|
| | M-SADA | AlpahFold2 |
| 1lafE | 0.998 | 0.998 |
| 1lagE | 0.998 | 0.998 |
| 1lahE | 0.998 | 0.998 |
| **2laoA** | 0.992 | 0.721 |
| **6mleE** | 0.998 | 0.998 |
| Average TM-score | 0.997 | 0.942 |

**Table S2.38** Multidomain protein cluster 38.

| PDB ID | Best TM-score | |
|---|---|---|
| | M-SADA | AlpahFold2 |
| **2xrnA** | 0.942 | 0.793 |
| 2xroA | 0.876 | 0.691 |
| **2xroE** | 0.876 | 0.691 |
| Average TM-score | 0.898 | 0.725 |

**Table S2.39** Multidomain protein cluster 39.

| PDB ID | Best TM-score | |
|---|---|---|
| | M-SADA | AlpahFold2 |
| **1m1gB** | 0.793 | 0.744 |
| **1nprA** | 0.715 | 0.732 |
| Average TM-score | 0.754 | 0.738 |

**Table S2.40** Multidomain protein cluster 40.

| PDB ID | Best TM-score | |
|---|---|---|
| | M-SADA | AlpahFold2 |
| **2g7uC** | 0.941 | 0.677 |
| **2g7uD** | 0.958 | 0.940 |
| Average TM-score | 0.950 | 0.809 |

**Table S2.41** Multidomain protein cluster 41.

| PDB ID | Best TM-score | |
|---|---|---|
| | M-SADA | AlpahFold2 |
| **4cbvA** | 0.979 | 0.971 |
| **4cbvB** | 0.935 | 0.531 |
| Average TM-score | 0.957 | 0.751 |

**Table S2.42** Multidomain protein cluster 42.

| PDB ID | Best TM-score | |
|---|---|---|
| | M-SADA | AlpahFold2 |
| 2yvyA | 0.966 | 0.961 |
| **2yvzA** | 0.957 | 0.522 |
| 2yvzB | 0.957 | 0.521 |
| 5x9gA | 0.961 | 0.969 |
| 5x9gB | 0.963 | 0.969 |
| **5x9gD** | 0.956 | 0.963 |
| Average TM-score | 0.960 | 0.817 |

**Table S2.43** Multidomain protein cluster 43.

| PDB ID | Best TM-score | |
|---|---|---|
| | M-SADA | AlpahFold2 |
| **4p0iB** | 0.988 | 0.689 |
| **5otcB** | 0.997 | 0.997 |
| Average TM-score | 0.992 | 0.843 |

**Table S2.44** Multidomain protein cluster 44.

| PDB ID | Best TM-score | |
|---|---|---|
| | M-SADA | AlpahFold2 |
| **4ehtB** | 0.985 | 0.967 |
| **4eiaA** | 0.962 | 0.746 |
| Average TM-score | 0.974 | 0.856 |

**Table S2.45** Multidomain protein cluster 45.

| PDB ID | Best TM-score | |
|---|---|---|
| | M-SADA | AlpahFold2 |
| 1sw1A | 0.992 | 0.990 |
| **1sw1B** | 0.992 | 0.991 |
| 1sw4A | 0.991 | 0.989 |
| 1sw4B | 0.992 | 0.990 |
| **1sw5A** | 0.974 | 0.984 |
| 1sw5B | 0.971 | 0.985 |
| 1sw5C | 0.972 | 0.988 |
| Average TM-score | 0.983 | 0.988 |

**Table S2.46** Multidomain protein cluster 46.

| PDB ID | Best TM-score | |
|---|---|---|
| | M-SADA | AlpahFold2 |
| 1klfB | 0.982 | 0.660 |
| 1klfD | 0.982 | 0.660 |
| 1klfF | 0.982 | 0.660 |
| 1klfH | 0.982 | 0.660 |
| 1klfJ | 0.968 | 0.657 |
| 1klfL | 0.968 | 0.657 |
| 1klfN | 0.968 | 0.657 |
| 1klfP | 0.968 | 0.657 |
| 1qunB | 0.976 | 0.660 |
| 1qunD | 0.976 | 0.660 |
| 1qunF | 0.976 | 0.660 |

| | | |
|---|---|---|
| 1qunH | 0.976 | 0.660 |
| 1qunJ | 0.947 | 0.654 |
| 1qunL | 0.947 | 0.660 |
| 1qunN | 0.947 | 0.654 |
| 1qunP | 0.947 | 0.660 |
| **4j3oH** | 0.907 | 0.800 |
| 4xo9A | 0.908 | 0.824 |
| 4xoaA | 0.911 | 0.822 |
| 4xoaC | 0.911 | 0.829 |
| 4xoaE | 0.913 | 0.829 |
| 4xoaG | 0.912 | 0.827 |
| 4xobA | 0.927 | 0.834 |
| 4xobC | 0.925 | 0.851 |
| 4xobE | 0.927 | 0.847 |
| **4xobG** | 0.987 | 0.593 |
| Average TM-score | 0.951 | 0.715 |

**Table S2.47** Multidomain protein cluster 47.

| PDB ID | Best TM-score | |
|---|---|---|
| | M-SADA | AlpahFold2 |
| 1p7hL | 0.941 | 0.622 |
| 1p7hM | 0.892 | 0.633 |
| **1p7hN** | 0.941 | 0.622 |
| **1p7hO** | 0.892 | 0.633 |
| Average TM-score | 0.917 | 0.628 |

**Table S2.48** Multidomain protein cluster 48.

| PDB ID | Best TM-score | |
|---|---|---|
| | M-SADA | AlpahFold2 |
| 3fxqA | 0.769 | 0.693 |
| 3fxqB | 0.684 | 0.980 |
| 3fxrA | 0.768 | 0.692 |
| 3fxrB | 0.687 | 0.970 |
| **3fxuA** | 0.766 | 0.691 |
| 3fzjA | 0.763 | 0.695 |
| 3fzjB | 0.684 | 0.978 |
| 3fzjC | 0.767 | 0.695 |
| 3fzjD | 0.684 | 0.974 |
| 3fzjE | 0.770 | 0.695 |
| 3fzjF | 0.684 | 0.976 |
| 3fzjG | 0.763 | 0.695 |
| **3fzjH** | 0.684 | 0.976 |
| 3fzjI | 0.764 | 0.694 |
| 3fzjJ | 0.684 | 0.977 |
| Average TM-score | 0.728 | 0.825 |

**Table S2.49** Multidomain protein cluster 49.

| PDB ID | Best TM-score | |
|---|---|---|
| | M-SADA | AlpahFold2 |
| **4fl4C** | 0.717 | 0.604 |
| **4fl4F** | 0.676 | 0.540 |
| Average TM-score | 0.696 | 0.572 |

**Table S2.50** Multidomain protein cluster 50.

| PDB ID | Best TM-score | |
|---|---|---|
| | M-SADA | AlpahFold2 |
| **1k23C** | 0.987 | 0.662 |
| **1wpmB** | 0.977 | 0.979 |
| 2hawB | 0.980 | 0.981 |
| Average TM-score | 0.981 | 0.874 |

**Table S2.51** Multidomain protein cluster 51.

| PDB ID | Best TM-score | |
|---|---|---|
| | M-SADA | AlpahFold2 |
| **3wfiB** | 0.983 | 0.759 |
| **5x20A** | 0.993 | 0.990 |
| 5x20C | 0.992 | 0.988 |
| Average TM-score | 0.989 | 0.912 |

**Table S2.52** Multidomain protein cluster 52.

| PDB ID | Best TM-score | |
|---|---|---|
| | M-SADA | AlpahFold2 |
| **4a5lA** | 0.995 | 0.730 |
| 4cbqA | 0.996 | 0.728 |
| 4ccqA | 0.994 | 0.730 |
| 4ccrB | 0.873 | 0.992 |
| **4ccrC** | 0.870 | 0.981 |
| Average TM-score | 0.946 | 0.832 |

**Table S2.53** Multidomain protein cluster 53.

| PDB ID | Best TM-score | |
|---|---|---|
| | M-SADA | AlpahFold2 |
| 1mwkA | 0.936 | 0.764 |
| **2qu4A** | 0.902 | 0.626 |
| 4a6jA | 0.991 | 0.986 |
| 4a6jB | 0.991 | 0.986 |
| 4a6jC | 0.991 | 0.986 |
| 4a6jD | 0.991 | 0.986 |
| 4a6jE | 0.991 | 0.986 |
| **4a6jF** | 0.991 | 0.986 |
| 4a6jG | 0.991 | 0.986 |
| 4a6jH | 0.991 | 0.986 |
| 4a6jI | 0.991 | 0.986 |
| 4a6jJ | 0.991 | 0.986 |
| Average TM-score | 0.979 | 0.938 |

**Table S2.54** Multidomain protein cluster 54.

| PDB ID | Best TM-score | |
|---|---|---|
| | M-SADA | AlpahFold2 |
| 1bpdA | 0.722 | 0.743 |
| 1huoA | 0.825 | 0.928 |
| 1huoB | 0.820 | 0.918 |
| 1huzA | 0.827 | 0.926 |
| 1huzB | 0.822 | 0.921 |

| | | |
|---|---|---|
| 2bpfA | 0.748 | 0.784 |
| **2bpgA** | 0.814 | 0.907 |
| 2bpgB | 0.811 | 0.895 |
| **3uxnA** | 0.719 | 0.745 |
| 3uxnB | 0.726 | 0.746 |
| Average TM-score | 0.783 | 0.851 |

**Table S2.55** Multidomain protein cluster 55.

| PDB ID | Best TM-score | |
|---|---|---|
| | M-SADA | AlpahFold2 |
| **1x2gC** | 0.823 | 0.679 |
| 1x2hC | 0.830 | 0.679 |
| 3a7aA | 0.971 | 0.972 |
| **3a7aC** | 0.973 | 0.973 |
| 3a7rA | 0.975 | 0.976 |
| Average TM-score | 0.915 | 0.856 |

**Table S2.56** Multidomain protein cluster 56.

| PDB ID | Best TM-score | |
|---|---|---|
| | M-SADA | AlpahFold2 |
| **1z15A** | 0.936 | 0.684 |
| **1z16A** | 0.998 | 0.995 |
| 1z17A | 0.997 | 0.995 |
| 1z18A | 0.995 | 0.993 |
| Average TM-score | 0.982 | 0.917 |

**Table S2.57** Multidomain protein cluster 57.

| PDB ID | Best TM-score | |
|---|---|---|
| | M-SADA | AlpahFold2 |
| **4oqoA** | 0.665 | 0.651 |
| 4oqoB | 0.671 | 0.652 |
| 5cryA | 0.985 | 0.984 |
| 5cryB | 0.986 | 0.984 |
| 5hbcA | 0.985 | 0.984 |
| 5hbcB | 0.986 | 0.984 |
| **7equA** | 0.985 | 0.984 |
| 7equB | 0.985 | 0.984 |
| Average TM-score | 0.906 | 0.901 |

**Table S2.58** Multidomain protein cluster 58.

| PDB ID | Best TM-score | |
|---|---|---|
| | M-SADA | AlpahFold2 |
| **2yglA** | 0.982 | 0.525 |
| **2ygmA** | 0.605 | 0.579 |
| Average TM-score | 0.793 | 0.552 |

**Table S2.59** Multidomain protein cluster 59.

| PDB ID | Best TM-score | |
|---|---|---|
| | M-SADA | AlpahFold2 |
| 2r6cA | 0.689 | 0.629 |
| 2r6cB | 0.799 | 0.605 |
| 2r6cC | 0.683 | 0.636 |
| 2r6cD | 0.758 | 0.603 |
| 2r6cE | 0.693 | 0.641 |
| 2r6cF | 0.795 | 0.603 |
| 4m4wA | 0.660 | 0.598 |
| 4m4wB | 0.739 | 0.574 |
| 4m4wC | 0.652 | 0.607 |
| 4m4wD | 0.727 | 0.582 |
| **4m4wE** | 0.660 | 0.614 |
| **4m4wF** | 0.762 | 0.568 |
| Average TM-score | 0.718 | 0.605 |

**Table S2.60** Multidomain protein cluster 60.

| PDB ID | Best TM-score | |
|---|---|---|
| | M-SADA | AlpahFold2 |
| **3dplC** | 0.829 | 0.979 |
| **3dqvD** | 0.942 | 0.796 |
| Average TM-score | 0.885 | 0.888 |

**Table S2.61** Multidomain protein cluster 61.

| PDB ID | Best TM-score | |
|---|---|---|
| | M-SADA | AlpahFold2 |
| 4kbfA | 0.651 | 0.650 |
| **4kbfB** | 0.678 | 0.587 |
| **4kbgB** | 0.744 | 0.703 |
| Average TM-score | 0.691 | 0.646 |

**Table S2.62** Multidomain protein cluster 62.

| PDB ID | Best TM-score | |
|---|---|---|
| | M-SADA | AlpahFold2 |
| **4by9C** | 0.653 | 0.374 |
| 4by9F | 0.738 | 0.493 |
| 4by9I | 0.651 | 0.379 |
| **4by9L** | 0.733 | 0.499 |
| Average TM-score | 0.694 | 0.436 |

**Table S2.63** Multidomain protein cluster 63.

| PDB ID | Best TM-score | |
|---|---|---|
| | M-SADA | AlpahFold2 |
| **4hh2A** | 0.829 | 0.657 |
| **4hh2C** | 0.711 | 0.697 |
| Average TM-score | 0.770 | 0.677 |

**Table S2.64** Multidomain protein cluster 64.

| PDB ID | Best TM-score | |
|---|---|---|
| | M-SADA | AlpahFold2 |
| 1anfA | 0.996 | 0.996 |
| 1dmbA | 0.869 | 0.837 |
| 1ez9A | 0.882 | 0.854 |
| 1ez9B | 0.857 | 0.820 |
| 1jw4A | 0.847 | 0.806 |
| 1jw5A | 0.846 | 0.806 |
| 1llsA | 0.845 | 0.804 |
| 1ompA | 0.854 | 0.816 |
| **2d21A** | 0.654 | 0.593 |
| 2mv0A | 0.859 | 0.809 |
| **2n44A** | 0.835 | 0.763 |
| 2n45A | 0.869 | 0.833 |
| 2r6gE | 0.875 | 0.844 |
| 3mbpA | 0.997 | 0.997 |
| 4mbpA | 0.996 | 0.997 |
| 5ldfM | 0.996 | 0.996 |
| 5ldfN | 0.996 | 0.996 |
| 5ldfO | 0.996 | 0.996 |
| 5ldfP | 0.996 | 0.996 |
| 5ldfQ | 0.996 | 0.996 |
| 5ldfR | 0.996 | 0.996 |
| 5ldfS | 0.996 | 0.996 |
| 5ldfT | 0.996 | 0.996 |
| 5ldfU | 0.996 | 0.996 |
| 5ldfV | 0.996 | 0.996 |
| 5ldfW | 0.996 | 0.996 |
| 5ldfX | 0.996 | 0.996 |
| Average TM-score | 0.927 | 0.908 |

**Table S2.65** Multidomain protein cluster 65.

| PDB ID | Best TM-score | |
|---|---|---|
| | M-SADA | AlpahFold2 |
| 4ct8A | 0.974 | 0.935 |
| **4ct9A** | 0.980 | 0.923 |
| 4ctaB | 0.975 | 0.997 |
| 4uocA | 0.976 | 0.933 |
| 4uuwA | 0.980 | 0.925 |
| **4uuwB** | 0.975 | 0.998 |
| Average TM-score | 0.976 | 0.952 |

**Table S2.66** Multidomain protein cluster 66.

| PDB ID | Best TM-score | |
|---|---|---|
| | M-SADA | AlpahFold2 |
| 1w26A | 0.860 | 0.946 |
| **1w26B** | 0.815 | 0.911 |
| 5owiA | 0.617 | 0.589 |
| 5owiB | 0.618 | 0.590 |
| **5owjA** | 0.563 | 0.572 |
| 5owjB | 0.562 | 0.571 |
| Average TM-score | 0.673 | 0.696 |

**Table S2.67** Multidomain protein cluster 67.

| PDB ID | Best TM-score | |
|---|---|---|
| | M-SADA | AlpahFold2 |
| 3zziA | 0.910 | 0.755 |
| 3zziD | 0.826 | 0.820 |
| 3zziE | 0.907 | 0.764 |
| 3zziF | 0.904 | 0.864 |
| **3zziH** | 0.885 | 0.834 |
| **4ab7A** | 0.983 | 0.731 |
| 4ab7D | 0.858 | 0.852 |
| 4ab7E | 0.942 | 0.858 |
| 4ab7F | 0.899 | 0.840 |
| 4ab7G | 0.877 | 0.874 |
| Average TM-score | 0.899 | 0.819 |

**Table S2.68** Multidomain protein cluster 68.

| PDB ID | Best TM-score | |
|---|---|---|
| | M-SADA | AlpahFold2 |
| 3vdxC | 0.956 | 0.825 |
| 4d9jC | 0.982 | 0.833 |
| 4d9jD | 0.925 | 0.923 |
| **4d9jE** | 0.986 | 0.884 |
| 4d9jF | 0.984 | 0.831 |
| 4d9jG | 0.956 | 0.836 |
| 4d9jI | 0.986 | 0.827 |
| **4d9jK** | 0.986 | 0.829 |
| Average TM-score | 0.970 | 0.848 |

**Table S2.69** Multidomain protein cluster 69.

| PDB ID | Best TM-score | |
|---|---|---|
| | M-SADA | AlpahFold2 |
| 4f0pB | 0.905 | 0.932 |
| 4f0pC | 0.944 | 0.682 |
| 4f0pD | 0.935 | 0.693 |
| 4f0qA | 0.906 | 0.930 |
| 4f0qB | 0.903 | 0.931 |
| 4f0qC | 0.937 | 0.691 |
| 4f0qD | 0.942 | 0.694 |
| 4r28A | 0.904 | 0.936 |
| **4r28B** | 0.905 | 0.928 |
| **4r28C** | 0.945 | 0.678 |
| 4r28D | 0.946 | 0.677 |
| Average TM-score | 0.925 | 0.797 |

**Table S2.70** Multidomain protein cluster 70.

| PDB ID | Best TM-score | |
|---|---|---|
| | M-SADA | AlpahFold2 |
| 1w25A | 0.707 | 0.571 |
| **1w25B** | 0.706 | 0.571 |
| **2v0nB** | 0.717 | 0.730 |
| Average TM-score | 0.710 | 0.624 |

**Table S2.71** Multidomain protein cluster 71.

| PDB ID | Best TM-score | |
|---|---|---|
| | M-SADA | AlpahFold2 |
| 1aonA | 0.942 | 0.742 |
| 1aonB | 0.942 | 0.742 |
| 1aonC | 0.941 | 0.740 |
| 1aonD | 0.944 | 0.743 |
| 1aonE | 0.944 | 0.743 |
| 1aonF | 0.945 | 0.742 |
| **1aonG** | 0.942 | 0.740 |
| 1aonH | 0.990 | 0.944 |
| 1aonI | 0.990 | 0.945 |
| 1aonJ | 0.989 | 0.946 |
| 1aonK | 0.990 | 0.947 |
| 1aonL | 0.991 | 0.946 |
| 1aonM | 0.990 | 0.945 |
| 1aonN | 0.989 | 0.943 |
| 1gruA | 0.942 | 0.742 |
| 1gruB | 0.942 | 0.742 |
| 1gruC | 0.941 | 0.740 |
| 1gruD | 0.944 | 0.743 |
| 1gruE | 0.944 | 0.743 |
| 1gruF | 0.945 | 0.742 |
| 1gruG | 0.942 | 0.740 |
| 1gruH | 0.921 | 0.924 |
| 1gruI | 0.921 | 0.924 |
| 1gruJ | 0.921 | 0.925 |
| 1gruK | 0.923 | 0.929 |
| 1gruL | 0.923 | 0.928 |
| 1gruM | 0.922 | 0.924 |
| 1gruN | 0.920 | 0.923 |
| 1xckA | 0.992 | 0.947 |
| 1xckB | 0.985 | 0.958 |
| 1xckC | 0.992 | 0.935 |
| 1xckD | 0.993 | 0.944 |
| 1xckE | 0.986 | 0.947 |
| **1xckF** | 0.992 | 0.932 |
| 1xckG | 0.994 | 0.947 |
| 1xckH | 0.992 | 0.935 |
| 1xckI | 0.993 | 0.943 |
| 1xckJ | 0.994 | 0.946 |
| 1xckK | 0.990 | 0.955 |
| 1xckL | 0.994 | 0.946 |
| 1xckM | 0.992 | 0.952 |
| 1xckN | 0.987 | 0.928 |
| 2c7cA | 0.947 | 0.737 |
| 2c7cB | 0.947 | 0.739 |
| 2c7cC | 0.956 | 0.744 |
| 2c7cD | 0.951 | 0.741 |
| 2c7cE | 0.954 | 0.743 |
| 2c7cF | 0.950 | 0.739 |
| 2c7cG | 0.954 | 0.736 |
| 2c7dA | 0.954 | 0.741 |
| 2c7dB | 0.954 | 0.742 |
| 2c7dC | 0.959 | 0.742 |
| 2c7dD | 0.953 | 0.742 |
| 2c7dE | 0.953 | 0.743 |
| 2c7dF | 0.952 | 0.738 |
| 2c7dG | 0.953 | 0.738 |
| 2cgtA | 0.950 | 0.748 |

| | | |
|---|---|---|
| 2cgtB | 0.950 | 0.748 |
| 2cgtC | 0.950 | 0.748 |
| 2cgtD | 0.950 | 0.748 |
| 2cgtE | 0.950 | 0.748 |
| 2cgtF | 0.950 | 0.748 |
| 2cgtG | 0.950 | 0.748 |
| 2nwcA | 0.988 | 0.953 |
| 2nwcB | 0.991 | 0.933 |
| 2nwcC | 0.994 | 0.941 |
| 2nwcD | 0.994 | 0.948 |
| 2nwcE | 0.993 | 0.954 |
| 2nwcF | 0.994 | 0.947 |
| 2nwcG | 0.995 | 0.949 |
| 2nwcH | 0.994 | 0.952 |
| 2nwcI | 0.992 | 0.950 |
| 2nwcJ | 0.992 | 0.948 |
| 2nwcK | 0.993 | 0.951 |
| 2nwcL | 0.991 | 0.937 |
| 2nwcM | 0.992 | 0.938 |
| 2nwcN | 0.994 | 0.946 |
| 3e76A | 0.992 | 0.950 |
| 3e76B | 0.990 | 0.952 |
| 3e76C | 0.990 | 0.950 |
| 3e76D | 0.994 | 0.949 |
| 3e76E | 0.989 | 0.945 |
| 3e76F | 0.993 | 0.950 |
| 3e76G | 0.993 | 0.948 |
| 3e76H | 0.994 | 0.948 |
| 3e76I | 0.992 | 0.948 |
| 3e76J | 0.989 | 0.946 |
| 3e76K | 0.989 | 0.951 |
| 3e76L | 0.990 | 0.951 |
| 3e76M | 0.985 | 0.949 |
| 3e76N | 0.987 | 0.944 |
| 1aonA | 0.942 | 0.742 |
| 1aonB | 0.942 | 0.742 |
| 1aonC | 0.941 | 0.740 |
| 1aonD | 0.944 | 0.743 |
| 1aonE | 0.944 | 0.743 |
| 1aonF | 0.945 | 0.742 |
| 1aonG | 0.942 | 0.740 |
| 1aonH | 0.990 | 0.944 |
| 1aonI | 0.990 | 0.945 |
| 1aonJ | 0.989 | 0.946 |
| 1aonK | 0.990 | 0.947 |
| 1aonL | 0.991 | 0.946 |
| 1aonM | 0.990 | 0.945 |
| 1aonN | 0.989 | 0.943 |
| 1gruA | 0.942 | 0.742 |
| 1gruB | 0.942 | 0.742 |
| 1gruC | 0.941 | 0.740 |
| 1gruD | 0.944 | 0.743 |
| 1gruE | 0.944 | 0.743 |
| 1gruF | 0.945 | 0.742 |
| 1gruG | 0.942 | 0.740 |
| 1gruH | 0.921 | 0.924 |
| 1gruI | 0.921 | 0.924 |
| 1gruJ | 0.921 | 0.925 |
| 1gruK | 0.923 | 0.929 |
| 1gruL | 0.923 | 0.928 |

| | | |
|---|---|---|
| 1gruM | 0.922 | 0.924 |
| 1gruN | 0.920 | 0.923 |
| 1xckA | 0.992 | 0.947 |
| 1xckB | 0.985 | 0.958 |
| 1xckC | 0.992 | 0.935 |
| 1xckD | 0.993 | 0.944 |
| 1xckE | 0.986 | 0.947 |
| 1xckF | 0.992 | 0.932 |
| 1xckG | 0.994 | 0.947 |
| 1xckH | 0.992 | 0.935 |
| 1xckI | 0.993 | 0.943 |
| 1xckJ | 0.994 | 0.946 |
| 1xckK | 0.990 | 0.955 |
| 1xckL | 0.994 | 0.946 |
| 1xckM | 0.992 | 0.952 |
| 1xckN | 0.987 | 0.928 |
| 2c7cA | 0.947 | 0.737 |
| 2c7cB | 0.947 | 0.739 |
| 2c7cC | 0.956 | 0.744 |
| 2c7cD | 0.951 | 0.741 |
| 2c7cE | 0.954 | 0.743 |
| 2c7cF | 0.950 | 0.739 |
| 2c7cG | 0.954 | 0.736 |
| 2c7dA | 0.954 | 0.741 |
| 2c7dB | 0.954 | 0.742 |
| 2c7dC | 0.959 | 0.742 |
| 2c7dD | 0.953 | 0.742 |
| 2c7dE | 0.953 | 0.743 |
| 2c7dF | 0.952 | 0.738 |
| 2c7dG | 0.953 | 0.738 |
| 2cgtA | 0.950 | 0.748 |
| 2cgtB | 0.950 | 0.748 |
| 2cgtC | 0.950 | 0.748 |
| 2cgtD | 0.950 | 0.748 |
| 2cgtE | 0.950 | 0.748 |
| 2cgtF | 0.950 | 0.748 |
| 2cgtG | 0.950 | 0.748 |
| 2nwcA | 0.988 | 0.953 |
| 2nwcB | 0.991 | 0.933 |
| 2nwcC | 0.994 | 0.941 |
| 2nwcD | 0.994 | 0.948 |
| 2nwcE | 0.993 | 0.954 |
| 2nwcF | 0.994 | 0.947 |
| 2nwcG | 0.995 | 0.949 |
| 2nwcH | 0.994 | 0.952 |
| 2nwcI | 0.992 | 0.950 |
| 2nwcJ | 0.992 | 0.948 |
| 2nwcK | 0.993 | 0.951 |
| 2nwcL | 0.991 | 0.937 |
| 2nwcM. | 0.992 | 0.938 |
| 2nwcN | 0.994 | 0.946 |
| 3e76A | 0.992 | 0.950 |
| 3e76B | 0.990 | 0.952 |
| 3e76C | 0.990 | 0.950 |
| 3e76D | 0.994 | 0.949 |
| 3e76E | 0.989 | 0.945 |
| 3e76F | 0.993 | 0.950 |
| 3e76G | 0.993 | 0.948 |
| 3e76H | 0.994 | 0.948 |
| 3e76I | 0.992 | 0.948 |

| | | |
|---|---|---|
| 3e76J | 0.989 | 0.946 |
| 3e76K | 0.989 | 0.951 |
| 3e76L | 0.990 | 0.951 |
| 3e76M | 0.985 | 0.949 |
| 3e76N | 0.987 | 0.944 |
| Average TM-score | 0.969 | 0.866 |

**Table S2.72** Multidomain protein cluster 72.

| PDB ID | Best TM-score | |
|---|---|---|
| | M-SADA | AlpahFold2 |
| **3oojB** | 0.747 | 0.636 |
| **3oojH** | 0.589 | 0.589 |
| Average TM-score | 0.668 | 0.612 |

**Table S3.** Summary of domain structure assembly by M-SADA, M-SADA-w/o-A and M-SADA-w/o-H on 296 test proteins. Here, domain models for assembly are experimentally solved domain structures. The results of M-SADA, M-SADA-w/o-A and M-SADA-w/o-H are the best models. #TM-score≥0.8 represents the number of models with TM-score ≥ 0.8.

| Domain | Method | Average TM-score | Median TM-score | #TM-score ≥ 0.8 |
|---|---|---|---|---|
| 2dom (137) | M-SADA-best | 0.90 | 0.95 | 109 |
| | M-SADA-w/o-A | 0.81 | 0.85 | 77 |
| | M-SADA-w/o-H | 0.83 | 0.89 | 83 |
| 3dom (62) | M-SADA-best | 0.81 | 0.83 | 36 |
| | M-SADA-w/o-A | 0.71 | 0.70 | 22 |
| | M-SADA-w/o-H | 0.72 | 0.68 | 23 |
| m4dom (33) | M-SADA-best | 0.72 | 0.76 | 15 |
| | M-SADA-w/o-A | 0.58 | 0.58 | 3 |
| | M-SADA-w/o-H | 0.59 | 0.55 | 5 |
| 2dis (64) | M-SADA-best | 0.94 | 0.97 | 61 |
| | M-SADA-w/o-A | 0.88 | 0.93 | 50 |
| | M-SADA-w/o-H | 0.87 | 0.92 | 47 |
| All (296) | M-SADA-best | 0.87 | 0.93 | 221 |
| | M-SADA-w/o-A | 0.78 | 0.81 | 152 |
| | M-SADA-w/o-H | 0.79 | 0.82 | 158 |

### Supplementary Texts

#### Text S1. The process of detecting homologous proteins of target sequence

The process of detecting homologous proteins of target sequence can be divided into four stages, which corresponds to the searching of four structure databases (MPDB, PDB, AlphaFold DB90 and AlphaFold DB) through HMMER program. In stage 1, jackHMMER from HMMER3.3.2 is used to search MPDB. At a given sequence identity cutoff, the top five proteins are selected as homologous templates used in M-SADA, where the length of the homologous template must be greater than 80% of the length of the target sequence. If sufficient homologous templates are not generated, stage 2 is performed. In stage 2, the same way is used to search PDB. If no homologous template satisfying the conditions is generated, stage 3 is performed. In stage 3, jackHMMER is used to search homologous protein models of target in AlphaFold DB as templates, and only searches for the proteins corresponding to models with an average pLDDT ≥90. Stage 4 is performed if available homologous templates from the previous stages is still less than five, where the whole AlphaFold DB is searched. The homologous proteins are detected by iterative search against MPDB, PDB, AlphaFold DB90 and AlphaFold DB with gradually relaxed e-values of $1e^{-30}$, $1e^{-10}$, $1e^{-6}$ and $1e^{-3}$ until the number of available homologous templates reach five.

#### Text S2. Energy functions for domain assembly

The $E_{\text{total}}^{p}$ and $E_{\text{total}}^{\text{hybrid}}$ are defined as follows:

$$E_{\text{total}}^{p} = w_1 E_{\text{clash}} + w_2 E_{\text{bd}} + w_3 E_{\text{dist}} + w_4 E_{\text{tpl}}^{p} \tag{S1}$$

$$E_{\text{total}}^{\text{hybrid}} = w_5 E_{\text{clash}} + w_6 E_{\text{bd}} + w_7 E_{\text{dist}}^{\text{hybrid}} + w_8 E_{\text{tpl}}^{\text{hybrid}} \tag{S2}$$

in which each energy function term is explained.

$E_{\text{clash}}$ is designed to eliminate steric clashes between domains, which is computed as follow:

$$E_{\text{clash}} = \sum_{n=1}^{N\text{-}1} \sum_{m=n+1}^{N} \sum_{i=1}^{Rdom_n} \sum_{j=1}^{Rdom_m} E_{\text{clash}}^{i,j} \tag{S3}$$

$$E_{\text{clash}}^{i,j}=\begin{cases}\frac{1}{d_{C_\alpha}^{i,j}}, & d_{C_\alpha}^{i,j} < d_{C_\alpha}^{\text{cut}} \\ 0, & \text{otherwise}\end{cases}$$

where $d_{C_\alpha}^{\text{cut}} = 3.75\text{Å}$, $N$ is the number of domain models, $Rdom_n$ and $Rdom_m$ represent the number of residues in the $n$-th and $m$-th domain, respectively. $d_{C_\alpha}^{i,j}$ is the distance between $C_\alpha$s of the residue $i$ and $j$ in the evaluated decoy.

$E_{\text{bd}}$ is the domain boundary energy, which is defined as follow:

$$E_{\text{bd}} = \sum_{n=1}^{N-1} (bd_{C_\alpha}^{n,n+1} - bd_{C_\alpha}^{\text{cut}})^2 \tag{S4}$$

where the $bd_{C_\alpha}^{\text{cut}} = 3.8\text{Å}$, and $bd_{C_\alpha}^{n,n+1}$ is the $C_\alpha$ atom distance between the C-terminal residue of $n$-th domain and N-terminal residue of the ($n$ + 1)-th domain in the evaluated decoy. In the discontinuous domain proteins, the discontinuous domain is split into multiple segments because of the insertion of the continuous domains. Therefore, for discontinuous domain proteins, these discontinuous segments are treated as domains to calculate the energy.

$E_{\text{dist}}$ is the inter-domain distance potential, which is deep-learning inter-domain geometric constraints. In this study, we use DeepIDDP[1], our recently developed inter-domain distance prediction network, to predict the inter-residue distance of full-chain.

The inter-domain distance potential is defined as follow:

$$E_{\text{dist}} = \frac{\sum_{n=1}^{N\text{-}1}\sum_{m=n+1}^{N}\sum_{i=1}^{Rdom_n}\sum_{j=1}^{Rdom_m} E_{\text{dist}}^{i,j}}{count} \tag{S5}$$

$$E_{\text{dist}}^{i,j} = \min\{|d_{\text{decoy}, C_\beta}^{i,j} - d_{\text{distmap}}^{i,j}|, 100\}$$

where $d_{\text{decoy}, C_\beta}^{i,j}$ is the distance between $C_\beta$ atom ($C_\alpha$ for glycine) of the residue pair ($i$, $j$) in the evaluated decoy, and $d_{\text{distmap}}^{i,j}$ is the distance of the residue pair ($i$, $j$) in the predicted inter-domain distance map. Here, the distance map is generated by taking the bin value of the maximum predicted probability for each residue pair ($i$, $j$), and only those residue pairs with maximum predicted probability ≥0.30 are used to calculate $E_{\text{dist}}^{i,j}$. *count* is the number of residue pairs used to calculate $E_{\text{dist}}^{i,j}$.

$E_{\text{tpl}}^{p}$ is the template-based restraint energy used in the $p$-th population, which is defined as follow:

$$E_{\text{tpl}}^{p} = \frac{\sum_{n=1}^{N\text{-}1}\sum_{m=n+1}^{N}\sum_{i=1}^{Rdom_n}\sum_{j=1}^{Rdom_m} E_{\text{tpl}}^{p,\,i,\,j}}{L} \quad \text{(S6)}$$

$$E_{\text{tpl}}^{p,\,i,\,j} = |d_{\text{decoy, C}_\alpha}^{i,j} \text{ - } d_{\text{tplmap}}^{p,\,i,\,j}|$$

where $d_{\text{decoy, C}_\alpha}^{i,j}$ is the distance between $C_\alpha$ of the residue pair $(i, j)$ in the evaluated decoy of the $p$-th population and the $d_{\text{tplmap}}^{p,\,i,\,j}$ is a value of the residue pair $(i, j)$ in the $p$-th template-based distance map. The $p$-th template-based distance map is generated by two steps: (1) the $p$-th template is selected to build template-based domain positions according to a sliding-window based procedure, which is described in **Text S4** and **Figure S3** and (2) according to the template-based domain positions, the distance of each residue pair $(i, j)$ between domains is calculated to generate the template-based distance map.

$E_{\text{dist}}^{\text{hybrid}}$ is the inter-domain distance potential used in hybrid population, which is defined as follow:

$$E_{\text{dist}}^{\text{hybrid}} = \sum_{n=1}^{N\text{-}1}\sum_{m=n+1}^{N}\sum_{i=1}^{Rdom_n}\sum_{j=1}^{Rdom_m} \frac{\log\left(\left(d_{\text{decoy, C}_\beta}^{i,j} - \mu_{i,j}\right)^2 + 1\right)}{\sigma_{i,j}} \quad \text{(S7)}$$

where $\mu_{i,j}$ and $\sigma_{i,j}$ are the mean and standard deviation obtained by Gaussian fitting of the residue pair $(i, j)$ distance distribution predicted by DeepIDDP, respectively.

$E_{\text{tpl}}^{\text{hybrid}}$ is the template-based restraint energy used in hybrid population, which integrates all selected templates to generate template-based restraint to guide domain assembly. $E_{\text{tpl}}^{\text{hybrid}}$ is defined as follow:

$$E_{\text{tpl}}^{\text{hybrid}} = \text{TMscore}_{\text{tpl}}^{\text{best}} \times E_{\text{tpl}}^{\text{best}} + \frac{(1 \text{ - } \text{TMscore}_{\text{tpl}}^{\text{best}})\sum_{p=1}^{P}\text{TMscore}_{\text{tpl}}^{p} \times E_{\text{tpl}}^{p}}{\sum_{p=1}^{P}\text{TMscore}_{\text{tpl}}^{p}} \quad \text{(SS8)}$$

where $\text{TMscore}_{\text{tpl}}^{p}$ is the highest average TM-score of the N/C-domains among all the positions of the $p$-th template, which is described in **Text S4** and **Figure S3**. $P$ is the number of templates used in M-SADA. $\text{TMscore}_{\text{tpl}}^{\text{best}}$ is the highest TM-score among $\text{TMscore}_{\text{tpl}}^{p}$s, and $E_{\text{tpl}}^{\text{best}}$ is the template-based restraint energy with best template-based domain position.

The weighting parameters in equation (S1) are set as $w_1 = 5.2$, $w_2 = 2.0$ and $w_4 = 0.50$, and

$w_3$ is set as equation (S9). If $\text{TMscore}_{\text{tpl}}^{p}$ is greater than or equal to 0.85, $w_4$ is set to 6.0. The weighting parameters in equation (S2) are set as $w_5$ = 5.2, $w_6$ = 2.0, $w_7$ = (1 - $\text{TMscore}_{\text{tpl}}^{\text{ave}}$) and $w_8$ = 0.13, where the $\text{TMscore}_{\text{tpl}}^{\text{ave}}$ is calculated by equation (S10). If the$\text{TMscore}_{\text{tpl}}^{\text{ave}}$ is greater than or equal to 0.82, $w_8$ is set to 1.5.

$$w_3 = \begin{cases} 10, & \text{TMscore}_{\text{tpl}}^{p} < 0.70 \\ 5\ , & 0.70 \le \text{TMscore}_{\text{tpl}}^{p} < 0.80 \\ 1\ , & 0.80 \le \text{TMscore}_{\text{tpl}}^{p} < 0.90 \\ 0.5, & 0.90 \le \text{TMscore}_{\text{tpl}}^{p} \end{cases} \tag{S9}$$

$$\text{TMscore}_{\text{tpl}}^{\text{ave}} = \frac{\sum_{p=1}^{P} \text{TMscore}_{\text{tpl}}^{p}}{P} \tag{S10}$$

The weighting parameters in Eq. (S1) and (S2) are determined by maximizing the TM-scores between the M-SADA models and the native structures, which are optimized through an improved differential evolution algorithm[2]. Details on the determination of the weighting parameters can be found in **Text S5**.

**Text S3. Multiple population-based evolutionary algorithm for domain assembly**

For a multi-domain protein with *N* domains, the solution of domain assembly can be represented as a (6 × *N*)-dimensional target vector, and the solution *S* can be represented as follow:

$$S = (x_1, y_1, z_1, \phi_1, \psi_1, \omega_1, \ldots x_N, y_N, z_N, \phi_N, \psi_N, \omega_N)$$

where $x_N, y_N, z_N$ represents the translation vector of the *N*-th domain and $\phi_N, \psi_N, \omega_N$ represents the three rotation angles of the *N*-th domain.

Under the guidance of different energy functions, a two-stage differential evolution algorithm is proposed to determine the potential solutions. The exploration stage aims to prevent the algorithm from getting stuck in local optima and generate multiple superior topology structures for the exploitation stage. Thus, the mutation strategy of slow convergence speed and strong exploration capability is used in this stage. In the exploitation stage, the explored superior solutions rapidly converged to the minimum. Therefore, on the basis of the superior solutions generated in the previous stage, we further adjusted the local position of these solutions to generate the optimal solution, and the mutation strategy with fast convergence speed was used at this stage. The multiple population-based evolutionary algorithm is described as follows:

**Initialization.** The number of populations is set according to the number of energy functions. In each population, the population size is set to *NP*. Then, the initial population should maximize the coverage of the entire search space by uniformly randomizing individuals within the search space constrained by the prescribed minimum and maximum parameter bounds $BD_{\text{explore}}^{\min} = (x_1^{\min}, y_1^{\min}, z_1^{\min}, \phi_1^{\min}, \psi_1^{\min}, \omega_1^{\min}, \ldots, x_N^{\min}, y_N^{\min}, z_N^{\min}, \phi_N^{\min}, \psi_N^{\min}, \omega_N^{\min})$ and $BD_{\text{explore}}^{\max} = (x_1^{\max}, y_1^{\max}, z_1^{\max}, \phi_1^{\max}, \psi_1^{\max}, \omega_1^{\max}, \ldots, x_N^{\max}, y_N^{\max}, z_N^{\max}, \phi_N^{\max}, \psi_N^{\max}, \omega_N^{\max})$, where $N$ is the number of domains.

In the $p$-th population $\{S_{p,1}, S_{p,2}, \ldots, S_{p,NP}\}$, each initial solution $S_{p,i}$ is generated as follow:

$$S_{p,i} = BD_{\text{explore}}^{\min} + rand[0, 1) \times (BD_{\text{explore}}^{\max} - BD_{\text{explore}}^{\min})$$

where $i = \{1, 2, \ldots, NP\}$ and $rand[0, 1)$ represents a uniformly distributed random variable within the range [0, 1).

In hybrid population, the size of hybrid population is set to *NHP*, and each initial solution $S_{\text{hybrid},i}$ in the hybrid population $\{S_{\text{hybrid},1}, S_{\text{hybrid},2}, \ldots, S_{\text{hybrid},NHP}\}$ is generated as follow:

$$S_{\text{hybrid},i} = BD_{\text{explore}}^{\min} + rand[0, 1) \times (BD_{\text{explore}}^{\max} - BD_{\text{explore}}^{\min})$$

where $i = \{1, 2, \ldots, NHP\}$.

The generation counter $g_1$ (for populations) and $g_2$ (for hybrid population) are set to 1. The new solutions are generated by mutation and crossover operation, and a selection operation is performed to determine whether old solutions can be replaced by new solutions in the population (or hybrid population). In exploration stage, the mutation, crossover and selection operations are performed as follows.

**Exploration stage:**

1. **Mutation operation.** The $i$-th solution of the $p$-th population ($S_{p,i}$) is set to the target solution ($S_{p,\text{target}}$), and performed the operation 1.1. The $i$-th solution of the hybrid population ($S_{\text{hybrid},i}$) is set to the target solution ($S_{\text{hybrid, target}}$), and performed the operation 1.2.

**1.1.** Three mutually different solutions are randomly selected in current $p$-th population, namely, $S_{p,\,rand1}$, $S_{p,\,rand2}$ and $S_{p,\,rand3}$. The mutant vector ($V_{p,\,i}$) is generated as follow:

$$V_{p,\,i} = S_{p,\,rand1} + F_{p,\,i} \times (S_{p,\,rand2} - S_{p,\,rand3})$$

where $F_{p,\,i}$ is scale factor of the $i$-th solution in the $p$-th population. If the value of $F_{p,\,i} \times (S_{p,\,rand2} - S_{p,\,rand3})$ exceeds the range between $BD_{\text{explore}}^{\text{min}}$ and $BD_{\text{explore}}^{\text{max}}$, we randomly and uniformly reinitialize them between $BD_{\text{explore}}^{\text{min}}$ and $BD_{\text{explore}}^{\text{max}}$.

**1.2.** Three mutually different solutions are randomly selected in current hybrid population, namely, $S_{\text{hybrid},\,rand1}$, $S_{\text{hybrid},\,rand2}$ and $S_{\text{hybrid},\,rand3}$. The mutant vector ($V_{\text{hybrid},\,i}$) is generated as follow:

$$V_{\text{hybrid},\,i} = S_{\text{hybrid},\,rand1} + F_{\text{hybrid},\,i} \times (S_{\text{hybrid},\,rand2} - S_{\text{hybrid},\,rand3})$$

where $F_{\text{hybrid},\,i}$ is scale factor of the $i$-th solution in the hybrid population. If the value of $F_{\text{hybrid},\,i} \times (S_{\text{hybrid},\,rand2} - S_{\text{hybrid},\,rand3})$ exceeds the range between $BD_{\text{explore}}^{\text{min}}$ and $BD_{\text{explore}}^{\text{max}}$, we randomly and uniformly reinitialize them between $BD_{\text{explore}}^{\text{min}}$ and $BD_{\text{explore}}^{\text{max}}$.

2. **Crossover operation.** After the mutation operation, crossover operation is applied to the target solution and its corresponding mutant vector to generate a trial solution. In the $p$-th population, a trial solution of the $i$-th solution ($U_{p,\,i}$) is generated by operation 2.1. In the hybrid population, a trial solution of the $i$-th solution ($U_{\text{hybrid},\,i}$) is generated by operation 2.2.

**2.1.** In the $p$-th population, each dimension $j$ of $U_{p,\,i}$ is generated according to a crossover rate of the $i$-th solution for the $p$-th population in exploration stage ($CR_{p,\,i}^{\text{explore}}$), which is defined as follow:

$$U_{p,\,i}^{j} = \begin{cases} V_{p,\,i}^{j}, \text{ if } (\,rand[0,1) < CR_{p,\,i}^{\text{explore}}\,) \text{ or } (\,j = j_{\text{rand}}\,) \\ S_{p,\,\text{target}}^{j}, \text{ otherwise} \end{cases}$$

where $j_{\text{rand}}$ is a randomly chosen integer in the range [1, 6×$N$]. The crossover rate controls the fraction of parameter values copied from the mutant vector.

**2.2.** In the hybrid population, each dimension $j$ of $U_{\text{hybrid},\,i}$ is generated according to a crossover rate of the $i$-th solution for hybrid population in exploration stage ($CR_{\text{hybrid},\,i}^{\text{explore}}$), which is defined as follow:

$$U_{\text{hybrid}, i}^{j} = \begin{cases} V_{\text{hybrid}, i}^{j}, \text{ if } (\, rand[0,1) < CR_{\text{hybrid}, i}^{\text{explore}}) \text{ or } (j = j_{\text{rand}}) \\ S_{\text{hybrid, target}}^{j}, \text{ otherwise} \end{cases}$$

3. **Selection operation.** The trial solution is first pre-evaluated by the energy ($w_1E_{\text{clash}} + w_2E_{\text{bd}}$). If the energy ($w_1E_{\text{clash}} + w_2E_{\text{bd}}$) of the trial solution is lower than that of the target solution or the energy ($w_1E_{\text{clash}} + w_2E_{\text{bd}}$) is less $0.25 \times N$, the trial solution is further evaluated by $E_{\text{total}}^{p}$ (or $E_{\text{total}}^{\text{hybrid}}$). Otherwise, the trial solution whether can be evaluated by $E_{\text{total}}^{p}$ (or $E_{\text{total}}^{\text{hybrid}}$) in accordance with the Boltzmann acceptance probability, which is defined as follow:

$$prob_{\text{Boltzmann}} = e^{\left(-\frac{(w_1E_{\text{clash}}(U_{\text{trial}}) + w_2E_{\text{bd}}(U_{\text{trial}})) - (w_1E_{\text{clash}}(S_{\text{target}}) + w_2E_{\text{bd}}(S_{\text{target}}))}{\beta}\right)}$$

if a random number between 0 and 1 is less than $prob_{\text{Boltzmann}}$, then the trial solution can be further evaluated by $E_{\text{total}}^{p}$ (or $E_{\text{total}}^{\text{hybrid}}$). Here, $U_{\text{trial}}$ and $S_{\text{target}}$ represent the trial solution and target solution, respectively. If the trial solution does not pass the pre-evaluation, the $i$-th solution in the $p$-th population (or hybrid population) is replaced by target solution ($S_{p,\text{ target}}$ or $S_{\text{hybrid, target}}$). Otherwise, the $i$-th solution in the $p$-th population (or hybrid population) is replaced by trial solution ($U_{p,\, i}$ or $U_{\text{hybrid},\, i}$) according to energy function ($E_{\text{total}}^{p}$ or $E_{\text{total}}^{\text{hybrid}}$). In the $p$-th population, the $i$-th solution is replaced according to operation 3.1. In the hybrid population, the $i$-th solution is replaced according to operation 3.2.

**3.1.** For the trial solution in the $p$-th population, the selection operation can be expressed as follow:

$$S_{p,\, i} = \begin{cases} U_{p,\, i}, \text{ if } E_{\text{total}}^{p}(U_{p,\, i}) \le E_{\text{total}}^{p}(S_{p,\text{ target}}) \\ S_{p,\text{ target}}, \text{ otherwise} \end{cases}$$

**3.2.** For the trial solution in the hybrid population, the selection operation can be expressed as follow:

$$S_{\text{hybrid},\, i} = \begin{cases} U_{\text{hybrid},\, i}, \text{ if } E_{\text{total}}^{\text{hybrid}}(U_{\text{hybrid},\, i}) \le E_{\text{total}}^{\text{hybrid}}(S_{\text{hybrid, target}}) \\ S_{\text{hybrid, target}}, \text{ otherwise} \end{cases}$$

4. **Iteration.** Mutation, crossover and selection operations in exploration stage are performed for all solutions in each population (including hybrid population). Then, steps 4.1 and 4.2 are performed.

**4.1.** The generation counter increases by 1 ($g_1 = g_1 + 1$ or $g_2 = g_2 + 1$). The exploitation stage is performed if the generation counter ($g_1$ and $g_2$) is greater than 0.7 × $G_{max}$. Otherwise, continue to execute steps 4.2 of exploration.

**4.2.** For each learning period ($LP$), the solutions between the different populations are exchanged. That is, the solutions between the different populations are exchanged if the remainder of the generation counter ($g_1$ and $g_2$) divided by $LP$ is 0. In the exchange process, the best $k$ solutions from each population are used to replace the worst $P \times k$ solutions in the hybrid population. Here, $P$ is the number of populations in addition to the hybrid population. Then, the worst $k$ solutions of each population are replaced with the best $k$ solutions of the hybrid population. Then, steps 1~4 of exploration stage are executed.

**Exploitation stage:**

In exploitation stage, the mutation operation, minimum and maximum parameter bounds and crossover rate are updated to exploit the potential solutions. In exploitation stage, the mutation, crossover and selection operations are performed as follows.

1. **Mutation operation.** The $i$-th solution of the $p$-th population ($S_{p,\,i}$) is set to the target solution ($S_{p,\,\text{target}}$), and operation 1.1 is performed. The $i$-th solution of the hybrid population ($S_{\text{hybrid},\,i}$) is set to the target solution ($S_{\text{hybrid},\,\text{target}}$), and operation 1.2 is performed.

**1.1.** Two mutually different solutions are randomly selected in current $p$-th population, namely, $S_{p,\,rand1}$ and $S_{p,\,rand2}$. The mutant vector ($V_{p,\,i}$) is generated as follow:

$$V_{p,\,i} = S_{p,\,\text{target}} + F_{p,\,i} \times (S_{p,\,rand1} - S_{p,\,rand2})$$

if the value of $F_{p,\,i} \times (S_{p,\,rand1} - S_{p,\,rand2})$ exceeds the range between $BD^{\min}_{\text{exploit}}$ and $BD^{\max}_{\text{exploit}}$, we randomly and uniformly reinitialize them between $BD^{\min}_{\text{exploit}}$ and $BD^{\max}_{\text{exploit}}$.

**1.2.** Two mutually different solutions are randomly selected in current hybrid population, namely, $S_{\text{hybrid},\,rand1}$ and $S_{\text{hybrid},\,rand2}$. The mutant vector ($V_{\text{hybrid},\,i}$) is generated as follow:

$$V_{\text{hybrid},\,i} = S_{\text{hybrid},\,\text{target}} + F_{\text{hybrid},\,i} \times (S_{\text{hybrid},\,rand1} - S_{\text{hybrid},\,rand2})$$

If the value of $F_{\text{hybrid},\,i} \times (S_{\text{hybrid},\,rand2} - S_{\text{hybrid},\,rand3})$ exceeds the range between $BD^{\min}_{\text{exploit}}$ and $BD^{\max}_{\text{exploit}}$, we randomly and uniformly reinitialize them between $BD^{\min}_{\text{exploit}}$

and $BD_{\text{exploit}}^{\max}$.

2. **Crossover operation.** After the mutation operation, crossover operation is applied to the target solution and its corresponding mutant vector to generate a trial solution. In the $p$-th population, a trial solution of the $i$-th solution ($U_{p,i}$) is generated by operation 2.1. In the hybrid population, a trial solution of the $i$-th solution ($U_{\text{hybrid},i}$) is generated by operation 2.2.

   **2.1.** In the $p$-th population, the crossover operation is defined as follows,

$$U_{p,i}^{j} = \begin{cases} V_{p,i}^{j}, \text{ if} (\, rand[0,1) < CR_{p,i}^{\text{exploit}}) \text{ or } (j = j_{\text{rand}}) \\ S_{p,\text{ target}}^{j}, \text{ otherwise} \end{cases}$$

   where $CR_{p,i}^{\text{exploit}}$ is a crossover rate of the $i$-th solution for the $p$-th population in exploitation stage.

   **2.2.** In the hybrid population, the crossover operation is defined as follows,

$$U_{\text{hybrid},i}^{j} = \begin{cases} V_{\text{hybrid},i}^{j}, \text{ if} (\, rand[0,1) < CR_{\text{hybrid},i}^{\text{exploit}}) \text{ or } (j = j_{\text{rand}}) \\ S_{\text{hybrid, target}}^{j}, \text{ otherwise} \end{cases}$$

   where $CR_{\text{hybrid},i}^{\text{exploit}}$ is a crossover rate of the $i$-th solution for hybrid population in exploitation stage.

3. **Selection operation.** The selection operation of exploitation stage is the same as exploration stage.

4. **Iteration.** Mutation, crossover and selection operations in exploitation stage are performed for all solutions in each population (including hybrid population). Then, steps 4.1 and 4.2 are performed.

   **4.1.** The generation counter increases by 1 ($g_1 = g_1 + 1$ and $g_2 = g_2 + 1$). The potential solutions are output if the generation counter ($g_1$ and $g_2$) is greater than $G_{\max}$. Otherwise, step 4.2 of exploitation stage is executed.

   **4.2.** The solutions between the different populations are exchanged if the remainder of the generation counter ($g_1$ and $g_2$) divided by $LP$ is 0. The exchange process is the same as exploration stage. Then, steps 2~4 of exploitation stage are executed.

5. **Output.** In each population, the solution with the lowest $E_{\text{total}}^{p}$ is selected. In the hybrid

population, three solutions are selected. The rules for the selection of solutions in the hybrid population are as follows: (1) the solution with the lowest $E_{\text{total}}^{\text{hybrid}}$ is selected, named $S_{\text{hybrid, best}}$, (2) the solution with the min cosine similarity to $S_{\text{hybrid, best}}$ is selected, name $S_{\text{hybrid, Mbest}}$ and (3) the solution with the min cosine similarity to $S_{\text{hybrid, Mbest}}$ is selected, named $S_{\text{hybrid, MMbest}}$. The $S_{\text{hybrid, Mbest}}$ and $S_{\text{hybrid, Mbest}}$ are selected as follows:

$$S_{\text{hybrid, Mbest}} = \min\{\frac{S_{\text{hybrid, best}} \cdot S_{\text{hybrid, }i}}{\|S_{\text{hybrid, best}}\| \, \|S_{\text{hybrid, }i}\|}, i = 1, 2, 3, \ldots, NHP\}$$

$$S_{\text{hybrid, MMbest}} = \min\{\frac{S_{\text{hybrid, Mbest}} \cdot S_{\text{hybrid, }i}}{\|S_{\text{hybrid, Mbest}}\| \, \|S_{\text{hybrid, }i}\|}, i = 1, 2, 3, \ldots, NHP\}$$

All selected solutions are ranked by DeepUMQA2[3] for output.

**Parameter setting:**

Population size:

$NP = 20$;

Hybrid population size:

$NHP = NP \times P$;

Maximum number of iterations:

$G_{\max} = 10000$;

Learning period:

$LP = 100$;

The prescribed minimum parameter bounds of exploration stage:

$BD_{\text{explore}}^{\min} = (-5, -5, -5, 0, 0, 0, \ldots, -5, -5, -5, 0, 0, 0)$;

The prescribed maximum parameter bounds of exploration stage:

$BD_{\text{explore}}^{\max} = (5, 5, 5, 2\pi, \pi, 2\pi, \ldots, 5, 5, 5, 2\pi, \pi, 2\pi)$;

Scale factor of exploration stage in the $i$-th solution:

$F_{p,i} = 0.1 + 0.9 \times rand(0, 1), i = 1, 2, 3, \ldots, NP; p = 1, 2, 3, \ldots, P$;

$F_{\text{hybrid}, i} = 0.1 + 0.9 \times rand(0, 1), i = 1, 2, 3, \ldots, NHP$;

Crossover rate of exploration stage in the *i*-th solution:

$CR_{p,\,i}^{\text{explore}} = rand(0, 1), i = 1, 2, 3, \ldots, NP; p = 1, 2, 3, \ldots, P;$

$CR_{\text{hybrid},\,i}^{\text{explore}} = rand(0, 1), i = 1, 2, 3, \ldots, NHP;$

The prescribed minimum parameter bounds of exploitation stage:

$BD_{\text{exploit}}^{\min} = (-BoE, -BoE, -BoE, 0, 0, 0, \ldots, -BoE, -BoE, -BoE, 0, 0, 0);$

$BoE = 1 - \text{TMscore}_{\text{tpl}}^{\text{ave}}$, where $\text{TMscore}_{\text{tpl}}^{\text{ave}}$ is defined in supplementary text S2.

The prescribed maximum parameter bounds of exploitation stage:

$BD_{\text{exploit}}^{\max} = (BoE, BoE, BoE, 0.5, 0.25, 0.5, \ldots, BoE, BoE, BoE, 0.5, 0.25, 0.5);$

Scale factor of exploitation stage:

$F_{p,\,i} = 0.5, i = 1, 2, 3, \ldots, NP; p = 1, 2, 3, \ldots, P;$

$F_{\text{hybrid},\,i} = 0.5, i = 1, 2, 3, \ldots, NHP;$

Crossover rate of exploitation stage:

$CR_{p,\,i}^{\text{exploit}} = 0.5, i = 1, 2, 3, \ldots, NP; p = 1, 2, 3, \ldots, P;$

$CR_{\text{hybrid},\,i}^{\text{exploit}} = 0.5, i = 1, 2, 3, \ldots, NHP;$

**Text S4. Template-based distance map generation by sliding-window based alignment**

Since the domain alignments are performed separately, the aligned regions of domains may be far away from each other. In this case, a sliding-window based procedure is employed to recreate domain alignments so that neighboring domains have the reasonable positions from the template. Taking a protein with 2 domains shown in **Figure S3** as an example, the N-terminal domain of the query is first superposed at the N-terminal of the template, and C-terminal domain is superposed at all the right-hand positions of the N-terminal domain along the template sequence with a maximum gap of 10 residues from N-terminal domain. Next, the superposition of N-terminal domain is shifted by one residue to the C-terminal of the template and redo the C-terminal superpositions. This procedure is repeated with the N-terminal domain sliding through all positions along the templates, where C-terminal domain is always on the right hand of the N-terminal domains. The alignment with the highest average TM-score of the N/C-domains among all the positions is finally selected for building domain positions based the template. According to the domain positions generated in previous step, the residue pairs distances between domains are calculated to generate template-based distance map.

**Text S5. The determination of weighting parameters for the force field**

The energy functions for domain assembly are defined in Eq. (S1) and Eq. (S2). The weighting parameters of the force field are determined by maximizing the TM-score between the M-SADA model and the native structure over a training set of 300 non-redundant proteins, where the weighting parameters are optimized through a differential evolution algorithm[2]. The 300 training multidomain proteins are randomly selected from a subset fetched from MPDB with the criteria (resolution: 0.0 to 3.0, number of domains: 2 to 10, sequence identity of each: 30%), and the sequence identity to the benchmark set is < 30%.

In the parameters training process, M-SADA was run from the initial parameters $\omega_1 = 1.0$, $\omega_2 = 1.0$, $\omega_3 =$ Eq. (S9), $\omega_4 = 1.0$, $\omega_5 = 1.0$, $\omega_6 = 1.0$, $\omega_7 = (1 - \mathrm{TMscore}_{\mathrm{tpl}}^{\mathrm{ave}})$, and $\omega_8 = 1.0$. The $\omega_3$ is piecewise constant function because the effect of the inter-residue distance potential needs to be reduced when the quality of templates is good. If the $\mathrm{TMscore}_{\mathrm{tpl}}^{p}$ is greater than or equal to a cutoff ($\omega_9$), $\omega_4$ is fixed to 6.0. If the$\mathrm{TMscore}_{\mathrm{tpl}}^{\mathrm{ave}}$ is greater than or equal to a cutoff ($\omega_{10}$), $w_8$ is set to 1.5.

A differential evolution algorithm is used to search an optimal solution ($\omega_1$, $\omega_2$, $\omega_4$, $\omega_5$, $\omega_6$, $\omega_8$, $\omega_9$, $\omega_{10}$) to maximize $f(\omega_1, \omega_2, \omega_4, \omega_5, \omega_6, \omega_8, \omega_9, \omega_{10})$ calculated with following equation:

$$f(\omega_1, \omega_2, \omega_4, \omega_5, \omega_6, \omega_8, \omega_9, \omega_{10}) = \sum_{n=1}^{N} TMscore(\omega_1, \omega_2, \omega_4, \omega_5, \omega_6, \omega_8, \omega_9, \omega_{10})_n$$

where $TMscore(\omega_1, \omega_2, \omega_4, \omega_5, \omega_6, \omega_8, \omega_9, \omega_{10})_n$ represents the TM-score of the $n$-th training protein model assembled by M-SADA with the parameters ($\omega_1$, $\omega_2$, $\omega_4$, $\omega_5$, $\omega_6$, $\omega_8$, $\omega_9$, $\omega_{10}$). As a result, the final weighting parameters ($\omega_1$, $\omega_2$, $\omega_4$, $\omega_5$, $\omega_6$, $\omega_8$, $\omega_9$, $\omega_{10}$) is (5.2, 2.0, 0.5, 5.2, 2.0, 0.13, 0.85, 0.82). $w_3$ of Eq. (S1) is set as Eq. (S9) and $w_7$ of Eq. (S2) is set as $(1 - \mathrm{TMscore}_{\mathrm{tpl}}^{\mathrm{ave}})$.
